# Enhanced MC1R-signalling and pH modulation facilitate melanogenesis within late endosomes of BLOC-1-deficient melanocytes

**DOI:** 10.1101/2024.07.08.602505

**Authors:** Philip S. Goff, Shyamal Patel, Tom Carter, Michael S. Marks, Elena V. Sviderskaya

## Abstract

Photoprotective melanins in the skin are synthesised by epidermal melanocytes within specialised lysosome-related organelles called melanosomes. Melanosomes coexist with lysosomes; thus, melanocytes employ specific trafficking machineries to ensure correct cargo delivery to either the endolysosomal system or maturing melanosomes. Mutations in some of the protein complexes required for melanogenic cargo delivery, such as biogenesis of lysosome-related organelles complex 1 (BLOC-1), result in hypopigmentation due to mistrafficking of cargo to endolysosomes. We show that hypopigmented BLOC-1-deficient melanocytes retain melanogenic capacity that can be enhanced by treatment with cAMP elevating agents despite the mislocalisation of melanogenic proteins. The melanin formed in BLOC-1-deficient melanocytes is not generated in melanosomes but rather within late endosomes/lysosomes to which some cargoes mislocalise. Although these organelles generally are acidic, a cohort of late endosomes/lysosomes have a sufficiently neutral pH to facilitate melanogenesis, perhaps due to mislocalised melanosomal transporters and melanogenic enzymes. Modulation of the pH of late endosomes/lysosomes by genetic manipulation or via treatment with lysosomotropic agents significantly enhances the melanin content of BLOC-1-deficient melanocytes. Our data suggest that upregulation of mistargeted cargoes can facilitate reprogramming of a subset of endolysosomes to generate some functions of lysosome-related organelles.

## Introduction

Lysosome-related organelles (LROs) are a diverse group of tissue-specific organelles that share features and components with organelles of the endolysosomal system (reviewed by Bowman, Bi-Karchin, Le, and Marks^1^). One such LRO, the melanosome, is found within melanocytes, the melanin producing cells that reside within the eye, hair follicles and the basal layer of the epidermis within the skin. Melanin produced by epidermal melanocytes is transferred to surrounding keratinocytes where it is arranged to form supranuclear caps that help protect against ultraviolet radiation^2^. Despite these photoprotective properties, melanin synthesis (melanogenesis) is an oxidative process that generates toxic intermediates^3,4^. Therefore, melanogenesis is confined within melanocytes to melanosomes to guard against these potentially cytotoxic effects.

Melanosomes mature through four morphologically distinct stages^5^. Stage I melanosomes are unpigmented acidic precursors that correspond to the vacuolar domains of early endosomes^6^. These organelles accumulate the pigment-cell specific pre-melanosome protein (PMEL) and undergo significant remodelling to the characteristic ellipsoid shape associated with melanosomes to form stage II melanosomes. This process is driven by the generation and elongation of amyloid fibrils formed following the proteolytic cleavage of PMEL^7,8^. These unpigmented stage II melanosomes mature to partially pigmented stage III melanosomes and fully pigmented stage IV melanosomes, which, under physiological conditions, are transferred to surrounding epidermal keratinocytes or retained intracellularly within the eye.

The maturation from stage II to stage III melanosomes requires the coordinated delivery of proteins that facilitate melanogenesis. These include the melanogenic proteins tyrosinase, tyrosinase-related protein 1 (TYRP1), and dopachrome tautomerase (DCT) that are directly involved in the chemical synthesis of melanin. Accessory proteins are also required such as ATP7A, which supplies the crucial melanogenic enzyme tyrosinase with its cofactor, copper^9^, and ion channels and transporters (e.g. OCA2 and SLC45A2) that raise luminal pH^10,11^ to near neutral and thus favour optimal tyrosinase activity^12,13^. The expression of most of these melanogenic proteins is regulated by melanocortin-1 receptor (MC1R) signalling. MC1R is a G-protein coupled receptor (GPCR), that, once bound by its physiological agonist, α-melanocyte stimulating hormone (α-MSH), increases melanogenic protein expression and melanogenesis via a cyclic adenosine monophosphate (cAMP)-dependent pathway^14–16^. As such, numerous studies have focused on activating the MC1R pathway to increase photoprotective melanin^17–21^.

Several LROs, including melanosomes, coexist with conventional endolysosomal organelles in their respective cell types^6^. Therefore, cells that harbour LROs (including melanocytes) have specialised trafficking mechanisms to distinguish between endolysosomal cargoes and those that are destined for maturing LROs. Genetic disorders that disrupt these mechanisms affect the biogenesis and/or function of these organelles and underlie multiple systemic consequences. One group of such disorders is the Hermansky-Pudlak syndromes (HPS), syndromic disorders characterised minimally by oculocutaneous albinism and bleeding diathesis^22^. To date, 11 subtypes of HPS have been identified (HPS-1 to -11) each caused by mutations in a different gene^23–32^. The HPS genes encode specific subunits of four distinct obligate protein complexes - Adaptor protein-3 (AP-3) and Biogenesis of lysosome-related organelles complex-1, -2 and -3 (BLOC-1, -2 and -3) that facilitate the delivery of cargoes to maturing LROs including melanosomes. Loss of function of any subunit of one of these complexes destabilises the other subunits of that complex.

The delivery of newly synthesised resident melanosomal proteins to maturing melanosomes is predominantly via early endosomes, facilitated in part by the HPS proteins. Disruption of any of the HPS complexes therefore impedes the delivery of specific melanogenic cargoes to melanosomes, resulting in a variable hypopigmentation phenotype. The majority of known melanogenic cargoes require BLOC-1 for exit from early endosomes toward maturing melanosomes^9,33–35^. BLOC-1 is a ubiquitously expressed protein complex comprised of eight subunits^27,36–40^ that form a flexible linear chain^41^. BLOC-1 facilitates cargo exit from early endosomes by the generation and elongation of recycling endosomal tubules in a process also involving adaptor protein 1 (AP-1) and the kinesin-3 microtubule motor heavy chain kinesin family member 13A (KIF13A)^42–44^.

Mutations in BLOC-1 subunits are responsible for HPS -7,-8, -9 and -11 and result in the accumulation of key melanogenic cargoes, such as ATP7A, TYRP1 and OCA2, within early endosomes^9,34,35,45^. A cohort of these cargoes have also been shown to leak into the degradative pathway^33,34^. A cohort of tyrosinase is also dependent on BLOC-1 for its delivery to melanosomes and accumulates largely in late endosomes and lysosomes in BLOC-1-deficient melanocytes^34,46^. Nevertheless, a substantial cohort of tyrosinase is present within melanosomes of BLOC-1-deficient melanocytes but is minimally active due at least in part to depletion of copper^9^. Although BLOC-1-deficient melanocytes from murine HPS models are severely hypopigmented, they are not completely devoid of melanin^47,48^. This suggests that despite the mislocalisation of several key melanogenic proteins, BLOC-1-deficient melanocytes retain at least some basal melanogenic capacity. Here, we show that enhanced MC1R-cAMP signalling and/or direct modulation of organelle pH of BLOC-1-deficient melanocytes increases melanin content but not within melanosomes. Instead, the melanin accumulates within late-endocytic organelles. The role of pH in regulating tyrosinase activity and the dependence of pH regulators on BLOC-1 for targeting to melanosomes may partially explain the hypopigmentation phenotype observed in forms of BLOC-1-deficient HPS and the ability of MC1R signalling to partially restore pigmentation in a remodelled late endosome.

## Results

### Prolonged MC1R/cAMP signalling enhances melanin content of BLOC-1-deficient melanocytes

Previous studies have used cAMP elevating agents to stimulate the MC1R signalling pathway and increase photoprotective eumelanin within the skin^19–21^. We asked whether modulation of this pathway can enhance melanogenesis in cells that mistraffic melanogenic cargoes such as BLOC-1-deficient melanocytes. To test this, we compared the melanin content of wild type (melan-a) and BLOC-1-deficient melanocytes (melan-pa, lacking the BLOC1S6 subunit of BLOC-1) that had been either unstimulated or cultured for 7 days with the cAMP elevating agents cholera toxin (CT), NDP-MSH (a synthetic analogue of α-MSH) or the adenylate cyclase activator forskolin.

Unstimulated wild type melanocytes were pigmented, and melanin content increased upon treatment with cAMP elevating agents as shown by bright field microscopy (Figure 1A) and validated by a quantitative melanin content assay (Figure 1C). These data are consistent with previous studies on the effects of enhanced MC1R signalling on melanocytes with competent trafficking machineries^20,21,49^.

**FIGURE 1.**
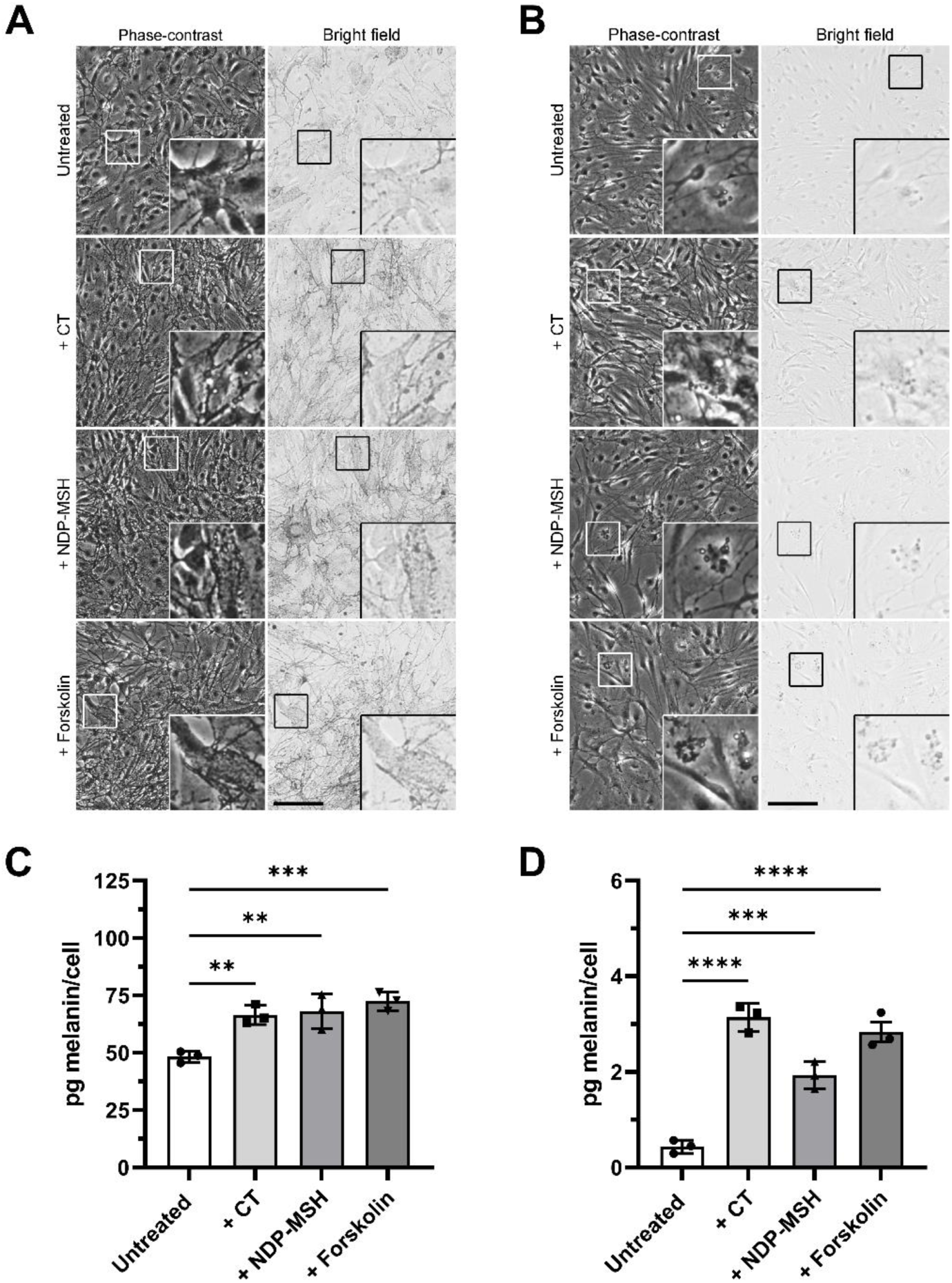
cAMP elevating agents increase melanin content of wild-type and BLOC-1-deficient melanocytes. **A, B.** Phase-contrast (left) and bright field (right) microscopy of wild-type melanocytes (melan-a, **A**) or BLOC-1-deficient melanocytes (melan-pa, **B**) either untreated or treated with 200 nM CT, 100 pM NDP-MSH or 20 µM forskolin for 7 days. Insets, boxed regions enlarged 3x. Scale bar, 100 µm. **C, D.** Quantification of melanin content of wild-type melan-a **(C)** or BLOC-1-deficient melan-pa **(D)** melanocytes cultured for 7 days in the presence or absence of either CT, NDP-MSH or forskolin as described above. Data presented show the means of 3 independent experiments and are expressed as mean pg melanin/cell ± SEM. Statistical significance relative to the untreated control was determined by one-way ANOVA with Dunnett’s multiple comparison test. **, p<0.01; ***, p<0.001; ****, p<0.0001.

In comparison, although unstimulated BLOC-1-deficient melanocytes were severely hypopigmented compared to wild type melanocytes at steady state as previously described^34^, treatment with cAMP elevating agents increased as visualised by bright field microscopy (Figure 1B) and confirmed by quantitative melanin content assay (Figure 1D). Similar results were obtained using another BLOC-1-deficient melanocyte line, melan-mu (lacking the BLOC1S5 subunit; Figure S1C), and the corresponding “rescued” line melan-mu:MuHA (BLOC-1^R^; Figure S1B). These data show that whilst melanin content of BLOC-1-deficient melanocytes is still much lower than that of wild type melanocytes, melanogenesis in BLOC-1-deficient melanocytes can be stimulated by cAMP elevating agents despite the lack of a critical transport route to melanosomes.

### Treatment with cAMP elevating agents enhances melanogenic protein expression in BLOC-1-deficient melanocytes

We next tested if increased melanogenic protein expression might contribute to the enhanced melanogenesis by cAMP elevating agents in BLOC-1-deficient melanocytes. Expression of the crucial melanogenic enzyme tyrosinase in wild type (melan-a), BLOC-1-deficient (melan-pa and melan-mu), and BLOC-1^R^ (melan-mu:MuHA) melanocytes treated for 7 days with cAMP elevating agents was quantified via western blotting (Figure 2A). Interestingly, significant increases in tyrosinase protein expression were only observed in BLOC-1-deficient melanocytes treated with cAMP elevating agents and not in wild type or BLOC-1^R^ melanocytes. The expression of other known melanogenic proteins (TYRP1 and PMEL) also showed a modest increase upon cAMP elevating agent treatment of BLOC-1-deficient melanocytes (Figure S2B and S2D) but not wild type and BLOC-1^R^ melanocytes (Figure S2A and S2C). The data suggest that cAMP elevating agent induced increases in melanin content of BLOC-1-deficient melanocytes may be at least in part due to enhanced expression of tyrosinase and perhaps other melanogenic proteins. The steady-state levels of the melanogenic protein TYRP1 are reduced in BLOC-1-deficient melanocytes^33^ likely because of partial mislocalisation to degradative organelles. Although tyrosinase primarily follows a BLOC-1-independent/AP-3-dependent pathway to melanosomes^46,50^, tyrosinase localisation to melanosomes is also reduced in BLOC-1-deficient melanocytes^34^. Accordingly, quantitative western blotting showed that steady-state levels of tyrosinase in BLOC-1-deficient (melan-mu and melan-pa) cells was reduced by ∼87% compared to wild type melan-a melanocytes (Figure 2B). This may result from mislocalisation and degradation of tyrosinase as has been previously reported for TYRP1^33,34^. Thus, the increased tyrosinase expression observed upon treatment with cAMP elevating agents might reflect both increased synthesis and reduced degradation.

**FIGURE 2.**
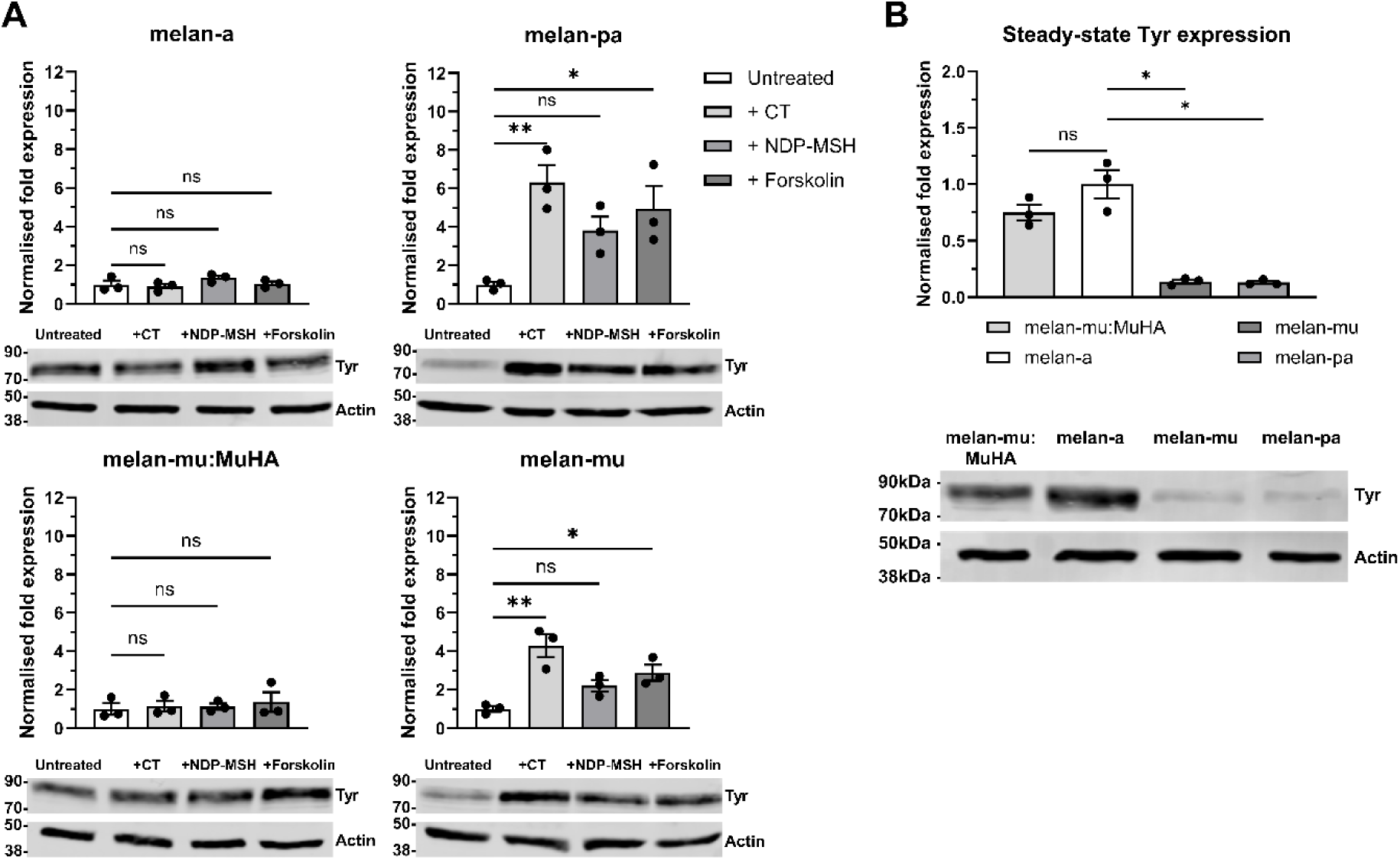
Tyrosinase protein expression in BLOC-1-deficient melanocytes is enhanced upon treatment with cAMP elevating agents. BLOC-1 competent (wild-type melan-a and BLOC-1^R^ melan-mu:MuHA) and BLOC-1-deficient (melan-pa and melan-mu) melanocytes were cultured in the absence or presence of the indicated cAMP elevating agents for 7 days, and then cell lysates were analysed by western blotting for tyrosinase (Tyr) and actin as a loading control. Representative blots are shown at the bottom, with positions of MW standards (in kDa) shown at the left, and quantifications from 3 independent biological replicates shown at top. Quantitative data are the mean normalised fold expression of tyrosinase protein relative to untreated control **(A)** or melan-a **(B)** ± SEM. Statistical significance relative to untreated control **(A)** or melan-a **(B)** tyrosinase expression was determined by one-way ANOVA with Dunnett’s multiple comparison test. Ns, not-significant; *, p<0.05; **, p<0.01.

### Melanin in BLOC-1-deficient melanocytes is predominantly formed within late-endosomal organelles

Cargoes destined for melanosomes primarily accumulate within early endosomes of BLOC-1-deficient melanocytes^9,34,35^, but a fraction of these proteins enter the degradative pathway^33^; for example, tyrosinase localisation to MVBs is increased nearly four-fold in BLOC-1-defcient melanocytes compared to BLOC-1^R^ melanocytes^34^. We therefore sought to assess in which compartments melanin accumulated in BLOC-1-deficient melanocytes.

Wild type and BLOC-1-deficient melanocytes were cultured with or without forskolin, and melanin (visualised by bright field microscopy) was localised relative to the mature melanosomal protein TYRP1 or the late endosomal/lysosomal membrane protein lysosome-associated membrane protein 2 (LAMP2) [visualised by immunofluorescence microscopy (IFM)]. Melanin within wild type melan-a (Figure 3A) and BLOC-1^R^ (data not shown but see Setty et al.^34^) melanocytes overlapped with TYRP1-positive organelles in both untreated and forskolin-treated cells (Figure 3A), and similar results were observed in melan-a cultured with other cAMP elevating agents (Figure S3). This indicates that regardless of treatment, melanin within wild type melanocytes accumulates as expected within melanosomes.

**FIGURE 3.**
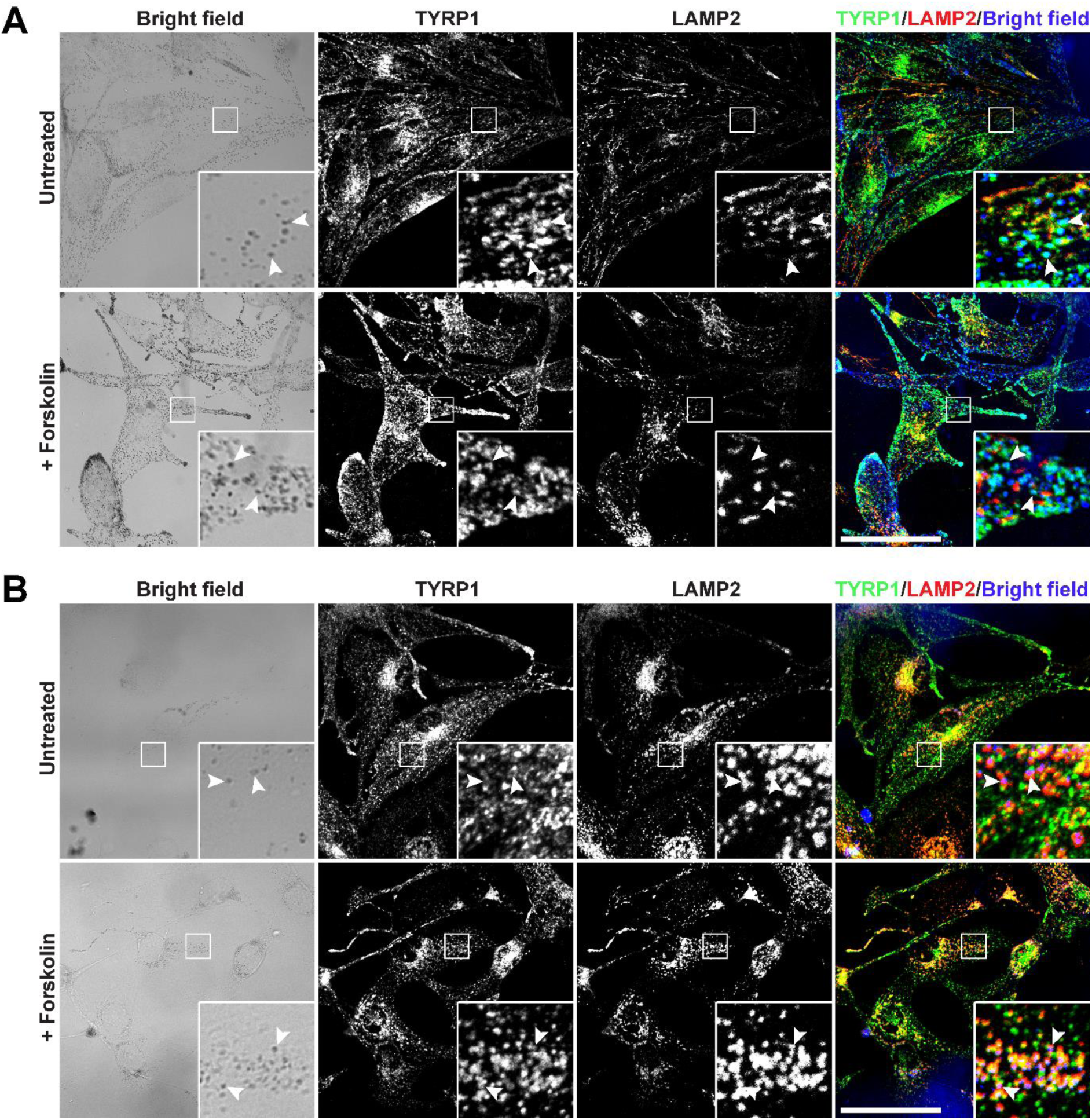
Melanin in BLOC-1-deficient melanocytes resides in LAMP2-positive organelles. Wild-type melan-a **(A)** or BLOC-1-deficient melan-pa **(B)** were treated or not with 20 µM forskolin for 7 days and then fixed, immunolabelled, and analysed by confocal IFM and bright field microscopy to determine the localisation of melanin (visible in bright-field images) in relation to melanosomes (TYRP1, green) and late endosomes/lysosomes (LAMP2, red). The bright-field images were inverted and pseudo-coloured blue in the merged image at the right. Insets, boxed regions enlarged 4.5x. Arrowheads show overlap of melanin with TYRP1 but not LAMP2 in melan-a **(A)** or with LAMP2 but not TYRP1 in melan-pa **(B)**. Scale bar, 50 µm.

In contrast, melanin within BLOC-1-deficient melan-pa melanocytes did not overlap with TYRP1-positive melanosomes and instead appeared to overlap with a small subset of LAMP2-positive structures irrespective of whether the cells were treated with cAMP elevating agents (Figure 3B and Figure S3). Similar results were seen in melan-mu, another BLOC-1-deficient melanocyte line (data not shown). The data suggest that melanin within BLOC-1-deficient melanocytes is formed not within bona-fide melanosomes but rather in late endosomes or lysosomes that harbour LAMP2.

LAMP2 and the related protein LAMP1 are highly abundant on lysosomal membranes^51,52^ but are also detected on late endosomes/multivesicular bodies (MVBs) and earlier endosomal compartments^53–55^. To better define where melanogenesis occurs in BLOC-1-deficient melanocytes and the nature of the LAMP2-containing organelles, we utilised both conventional transmission electron microscopy (TEM) and correlative light and electron microscopy (CLEM) analyses of the BLOC-1-deficient cell line melan-mu and BLOC-1^R^ melan-mu:MuHA with or without forskolin treatment.

By TEM, BLOC-1^R^ melanocytes contained numerous melanosomes of all stages (stage I – stage IV) under control conditions (Figure 4A) and upon forskolin treatment (Figure 4B). The morphology of stage III and IV melanosomes in BLOC-1^R^ melanocytes was typical of these organelles, i.e. predominantly ellipsoid in shape with striations visible within partially pigmented stage III organelles (Figure 4A’ and Figure 4B’) and fully occluded within stage IV melanosomes (Figure 4A’’ and Figure 4B’’). The mean length of the pigmented melanosomes in both untreated (428 ± 14 nm) and forskolin-treated (477 ± 17 nm) BLOC-1^R^ melanocytes (Figure 4G) is consistent with previous estimates of melanosomal length of approximately 500 nm^56^. In addition, the mean ratio of length to width is 1.77 ± 0.06 (untreated) and 1.97 ± 0.06 (forskolin-treated), consistent with the observed ellipsoid shape of these organelles.

**FIGURE 4.**
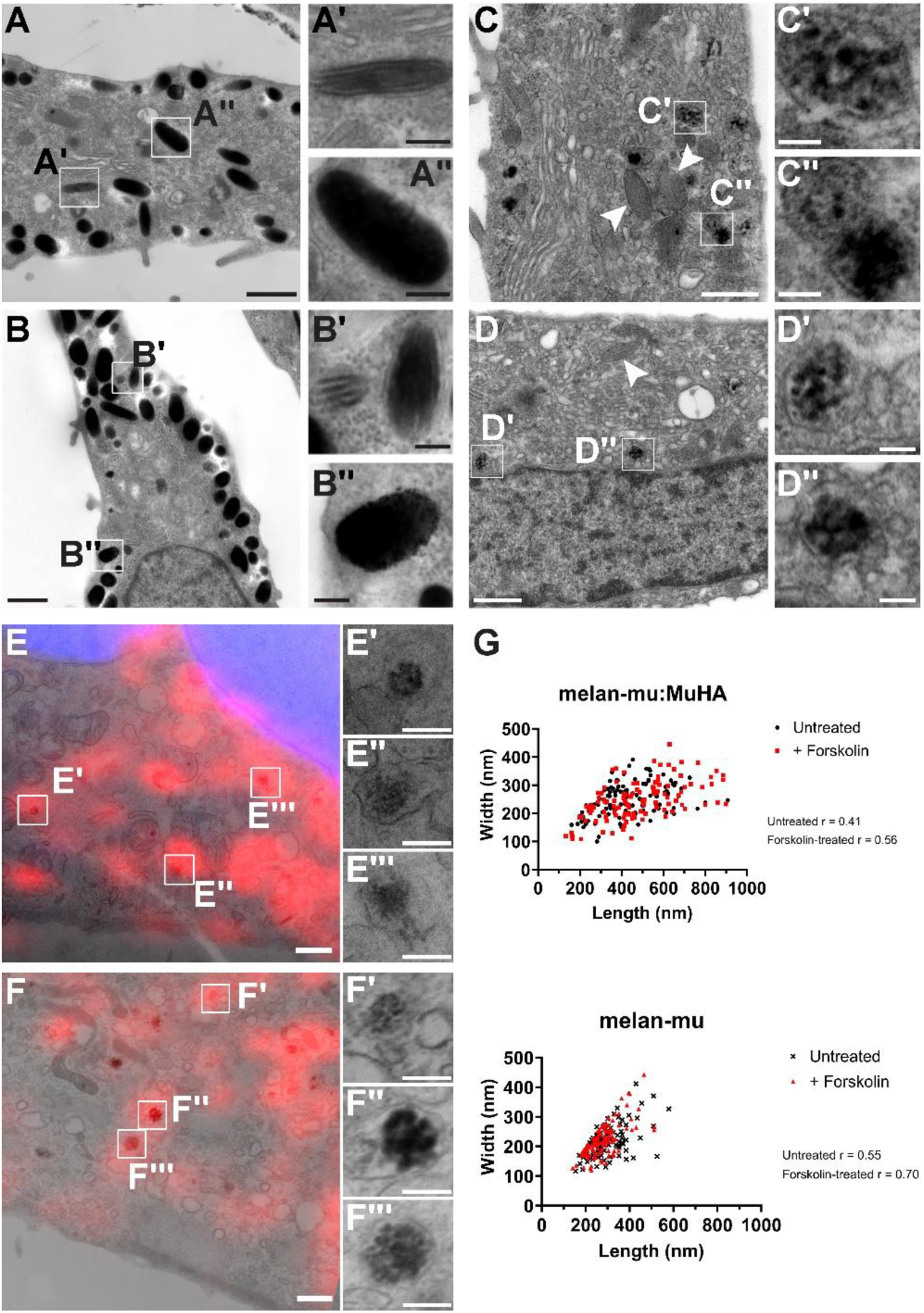
BLOC-1-deficient melanocytes produce melanin in LAMP2-positive organelles that resemble MVBs. Conventional electron microscopy analysis of untreated **(A)** or forskolin-treated **(B)** BLOC-1^R^ melanocytes (melan-mu:MuHA) or untreated **(C)** or forskolin-treated **(D)** BLOC-1-deficient melanocytes (melan-mu). Insets show 3x enlarged images of boxed regions in main panels. Arrowheads in **(C)** and **(D)** indicate unpigmented ellipsoid melanosomes. **A’**, a partially pigmented stage III melanosome; **A’’**, a fully-pigmented stage IV melanosome; **B’**, a stage III and stage IV melanosome; **B’’**, a fully-pigmented stage IV melanosome; **C’, C’’, D’** and **D’’**, circular organelles containing vesicular melanin deposits. **E-G.** CLEM analysis of untreated **(E)** or forskolin-treated **(F)** BLOC-1-deficient melanocytes (melan-mu) immunolabelled for LAMP2 (red) with DAPI staining the nucleus (blue). Insets show 4x enlarged images of boxed regions in main panels. **E’, E’’** and **E’’’**, ultrastructure of areas that are positive for LAMP2 in **E; F’, F’’** and **F’’’**, ultrastructure of areas that are positive for LAMP2 in **F**. **G**, Quantification of length and width of pigmented organelles in BLOC-1^R^ (melan-mu:MuHA) and BLOC-1-deficient (melan-mu) melanocytes cultured for 7 days with or without 20 µM forskolin. The length and width of a minimum of 100 pigmented organelles were obtained for each condition from electron microscopy images. No distinctions were made between different stages of melanosome maturation of if the organelle was a bona-fide melanosome. Scale bar in main panels, 800 nm. Scale bar insets, 200 nm.

In contrast to the BLOC-1^R^ cells, TEM analysis of BLOC-1-deficient melanocytes showed many fewer pigmented structures, consistent with the melanin quantification in Figure 1D and Figure S1C. As inferred from the IFM data, the few melanin deposits observed in either untreated (Figure 4C) or forskolin-treated (Figure 4D) BLOC-1-deficient melanocytes were not detected in bona-fide ellipsoid and striated structures, which were abundant in these cells but not pigmented as previously described^9,34^. Rather, the pigment-containing organelles (Figure 4C, 4D, 4E and 4F and accompanying insets) contained discrete clusters of melanin that appeared to surround and engulf intraluminal vesicles (ILVs) within MVBs. These organelles lacked the typical PMEL fibrils that are characteristic of stage II-IV melanosomes, were smaller in length than bona-fide melanosomes (301 ± 8 nm for untreated and 280 ± 7 nm for forskolin-treated), and had a more circular morphology (length to width ratio of 1.41 ± 0.04 untreated and 1.29 ± 0.03 for forskolin-treated; Figure 4G). Thus, melanin is formed in BLOC-1-deficient cells in organelles with morphological characteristics of late endosomes and not in traditional melanosomes.

We further analysed untreated and forskolin-treated BLOC-1-deficient melanocytes by CLEM. LAMP2-containing compartments with melanin pigments were identified by IFM and bright field microscopy in fixed cells on dishes with a raised alphanumeric coverslip on the bottom, and then cells were processed for conventional TEM. Corresponding regions were identified and aligned with the IFM images. CLEM confirmed that the melanin-containing organelles in BLOC-1-deficient melanocytes were positive for LAMP2 and were ultra-structurally similar to MVBs rather than ellipsoid and striated melanosomes (Figure 4E and 4F). CLEM also confirmed that many LAMP2-containing structures lacked pigment. Taken together, these data show that melanin within BLOC-1-deficient melanocytes is formed within a subset of late endocytic compartments that closely resemble MVBs, and is likely formed by mislocalised tyrosinase and other melanosomal components in these compartments due to the loss of BLOC-1.

### A subpopulation of late endosomes/lysosomes of BLOC-1-deficient melanocytes has elevated pH

Tyrosinase is the rate-limiting enzyme and is absolutely necessary for melanogenesis to occur. In order for optimal activity tyrosinase must be in a near neutral environment^12,13,57^; tyrosinase activity is abolished at pH <5.5, levels that are typically reported for late endosomes and lysosomes. The detection of melanin in late endosomes (based on morphology) of BLOC-1-deficient melanocytes suggests that a subset of these typically acidic organelles must be at least partially neutralised and that this subset should be expanded upon treatment with cAMP elevating agents.

To test this, we determined the pH of late endosomes/lysosomes in melanocytes by exploiting uptake of a fluid-phase cargo, dextran, conjugated to Oregon Green, a pH-dependent fluorophore. Cells were allowed to internalise Oregon Green-Dextran (OG-D) for 16 hours followed by a 2 hour chase into late endosomes and lysosomes before analysis by fluorometry or imaging by wide-field fluorescent microscopy. The pH-dependent fluorescence of internalised OG-D was confirmed by fluorometry upon exposure to previously described calibration solutions^58,59^ both in a cell-free environment and in wild type melanocytes (pKAs of 4.78 and 4.90 respectively; Figure S4A). Using the labelling protocol above (also see Methods), internalised OG-D in wild-type melanocytes overlapped with expressed LAMP1-tdTomato when analysed by wide-field fluorescent microscopy (Figure S4B), confirming that we were measuring the pH of late endosomes/lysosomes.

To measure lysosomal pH at steady-state and in response to treatment with cAMP elevating agents, wild type, BLOC-1^R^ and BLOC-1-deficient cell lines were cultured in the presence or absence of cAMP elevating agents before labelling with OG-D as described above. Fluorometry was used to measure both the resting fluorescence and the maximal fluorescence observed upon cell exposure to ammonium chloride. Lysosomal pH was calculated using a previously described equation^58^ also see Methods) using the parameters obtained from *in vivo* calibration curves (Figure S4A). The data showed that in all melanocyte lines tested the steady state pH of lysosomes is acidic (between pH 4.5 and 4.7), and treatment with cAMP elevating agents did not detectably neutralise pH at the population level (Figure 5A). However, since previous reports have shown that lysosomal pH is heterogeneous^60,61^, we investigated the subcellular heterogeneity of late endosomes/lysosomes using high magnification wide-field fluorescence microscopy in BLOC-1-deficient melanocytes that had internalised OG-D as described above and quantified the pH of individual late endosomes/lysosomes.

**FIGURE 5.**
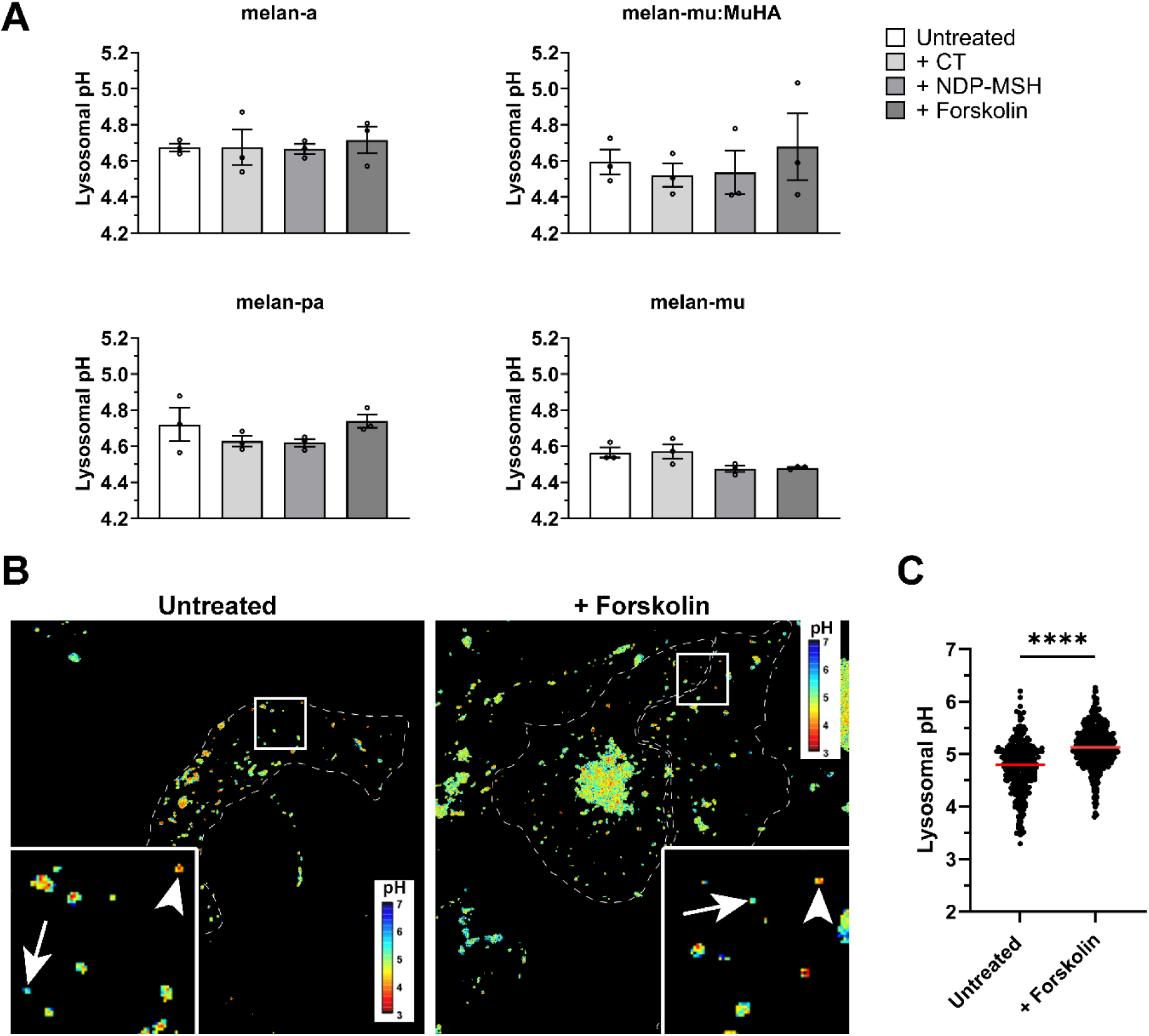
A subpopulation of late endosomes/lysosomes of BLOC-1-deficient melanocytes has elevated pH. BLOC-1 competent (melan-a and melan-mu:MuHA) and BLOC-1-deficient (melan-mu and melan-pa) melanocytes, cultured in the presence or absence of cAMP elevating agents for 7 days, were exposed to Oregon Green-Dextran overnight and chased for two hours to label late endosomes and lysosomes. **A.** Organellar pH was determined by fluorometry based on resting fluorescence relative to maximal fluorescence observed after exposure to ammonium chloride. Data represent mean ± SEM from 3 independent experiments. For each cell line, statistical analysis was performed by one-way ANOVA. No statistically significant differences were observed. **B.** Organellar fluorescence was assessed by live cell imaging before and after exposure to ammonium chloride and quantified in **C.** Shown are pseudo-coloured representations (note legend for scale) of the calculated lysosomal pH in untreated and forskolin-treated BLOC-1-deficient melanocytes. Arrows indicate lysosomes with a more neutral pH, and arrowheads indicate more acidic organelles. Insets show 3.5x enlarged image of boxed regions. Organellar pH was quantified from pseudo-coloured representations. The pH of 309 (untreated) and 369 (forskolin-treated) individual organelles was quantified from 4 and 5 cells respectively across 3 independent experiments. Red line represents mean pH of organelles for each condition. Statistical analysis was determined by two-tailed Student’s t-test. ****, p<0.0001.

As expected, most OG-D-containing late endosomes/lysosomes were acidic (arrowheads, Figure 5B and Figure 5C for quantification). However, a subset of organelles appeared to be more neutral with a pH closer to 6 (arrows Figure 5B and Figure 5C). Interestingly, the data showed that the mean intraluminal pH of late endosomes/lysosomes under control conditions is 4.7, and this increased to 5.1 upon treatment with the cAMP elevating agent forskolin. This increase in mean luminal pH upon forskolin treatment was not observed at the population level (Figure 5A). This difference in response to forskolin treatment may be due to a bias towards measuring well-isolated peripheral OG-D-containing late endosomes/lysosomes rather than those within the perinuclear region where we were unable to detect individual structures. Taken together the data show that the luminal pH of a subset of late endosomes/lysosomes in BLOC-1-deficient melanocytes is more neutral and can be further neutralised with forskolin-treatment. These structures are therefore likely to facilitate tyrosinase activity and subsequent melanogenesis.

### Genetic modulation of organelle pH is sufficient to enhance melanin outside of melanosomes

For melanogenesis to occur within the late endosomes of BLOC-1-deficient melanocytes, not only tyrosinase but also other melanogenic proteins, including regulators of pH such as the chloride ion channel OCA2^10^ and the putative proton/sugar transporter SLC45A2^62^, must be present to enable tyrosinase activity. Both OCA2 and SLC45A2 facilitate melanosome neutralisation to promote melanogenesis^10,11^ and at least OCA2 is targeted to melanosomes in a BLOC-1-dependent manner^35^. We therefore asked whether the overexpression of these regulators of organelle pH in BLOC-1-deficient melanocytes would partially neutralise the luminal pH of late endosomes and enhance melanogenesis in these organelles.

To test this we overexpressed HA-epitope tagged forms of OCA2^63^ or SLC45A2^11^ in BLOC-1-deficient melanocytes, and assayed for pigmentation by bright field microscopy and for lysosomal localisation by IFM relative to LAMP2. The data (Figure 6A; pigmentation quantified in Figure 6B) show that the overexpression of HA-OCA2 but not of HA-SLC45A2 enhances melanin content in BLOC-1-deficient melanocytes. Moreover, labelling for the HA-OCA2 transgene colocalised with both melanin and LAMP2 (Figure 6A). A similar increase in pigmentation in LAMP2-positive structures was seen upon overexpression of HA-OCA2 in the BLOC-1-deficient line melan-pa (Figure S5A; pigmentation quantified in Figure S5C). By contrast, overexpression of HA-OCA2 in hypopigmented *Oca2*-null melanocytes or of HA-SLC45A2 in *Slc45a2*-null melanocytes resulted in rescue of pigmentation (Figure S5C) but in structures that did not overlap with labelling for LAMP2 (Figure S5B), suggesting correct localisation to melanosomes as previously documented^11,63^. These data indicate that overexpression of OCA2 in BLOC-1-deficient melanocytes results in OCA2 delivery to a cohort of late endosomes/lysosomes where it enhances melanin content. Since OCA2 and SLC45A2 are targets of MITF and hence upregulated by MC1R signalling^64–66^, this may in part explain the effect of MC1R stimulation on enhancing pigmentation in cAMP elevating agent treated melanocytes.

**FIGURE 6.**
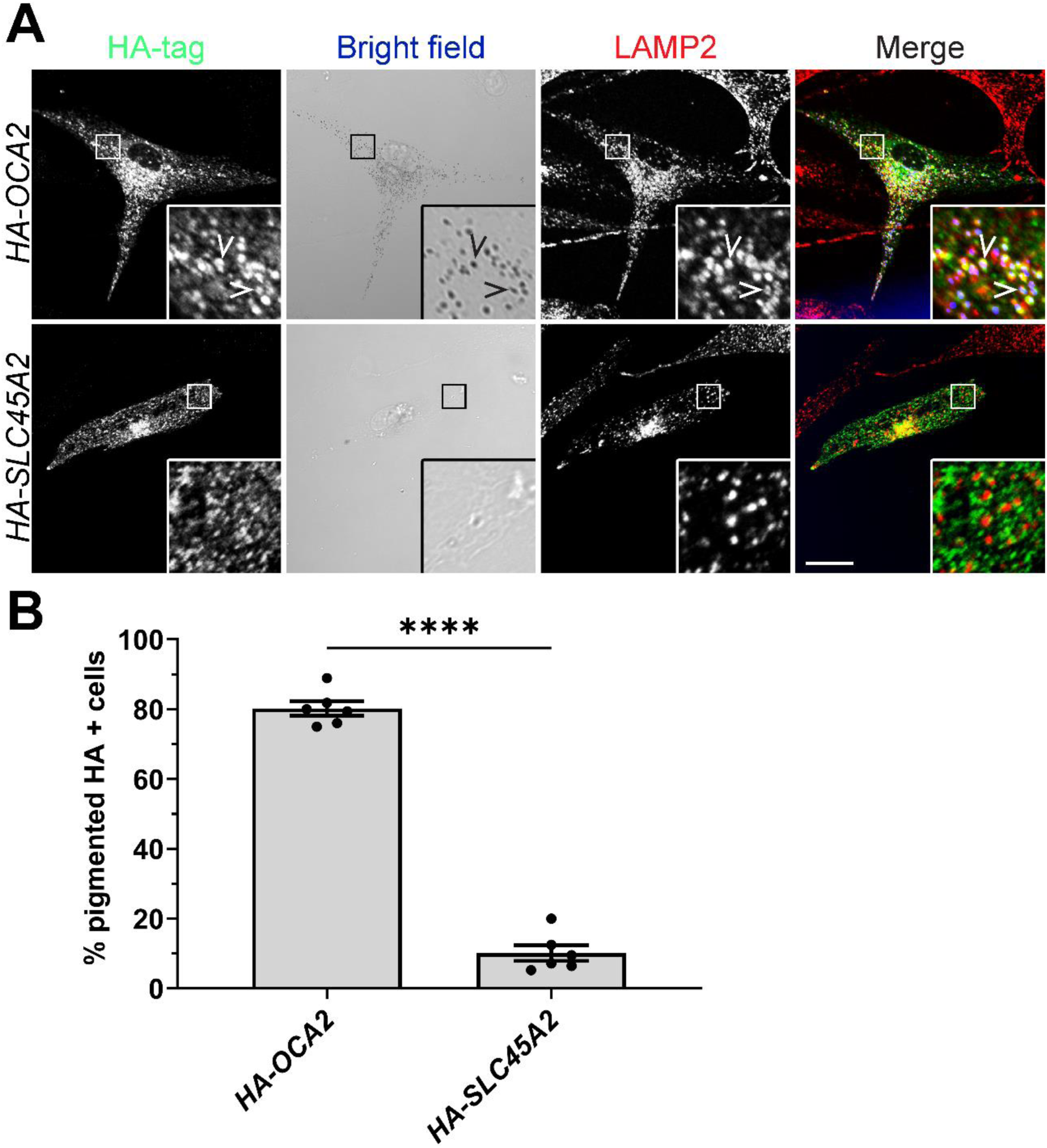
Overexpression of OCA2 in BLOC-1-deficient melanocytes partially restores pigmentation. BLOC-1-deficient melan-mu cells were transiently transfected with *HA-OCA2* or *HA-SLC45A2*, fixed and immunolabelled for HA and LAMP2, and analysed by confocal IFM and bright field microscopy to visualise melanin. **A.** Shown are each individual channel and an overlay on the right in which pigment granules are pseudo-coloured blue. Arrowheads indicate overlap between HA tag, LAMP2 and melanin in cells overexpressing *HA-OCA2*. Insets, boxed regions enlarged 4.5x. Scale bar, 20 µm. **B.** Quantification of the percentage of HA positive cells that were pigmented. Data represent mean ± SEM from 6 independent experiments. Sample size: 182 SLC45A2 HA positive cells and 194 OCA2 HA positive cells. Statistical significance relative to the % pigmented SLC45A2 HA positive cells was determined by two-tailed Student’s t-test. ****, p<0.0001.

### Direct modulation of organelle pH rapidly increases melanin within late endosomes/lysosomes

Numerous studies have reported that deacidification agents can significantly enhance melanogenesis in tyrosinase-positive amelanotic melanocytes^12,67–71^. To further test the effects of direct modulation of organelle pH on the melanin content of BLOC-1-deficient melanocytes, we assessed the effect on pigmentation of treatment with Bafilomycin A1 (BafA1), an inhibitor of the vacuolar ATPase that acidifies all post-Golgi compartments and methionine methyl ester (MME), a methylated amino acid that neutralises esterase-containing organelles such as lysosomes^72^.

BLOC-1-deficient melan-mu cells were treated for 2 or 16 hours with BafA1 or MME and visualised by bright-field microscopy (Figure 7A). Treatment with BafA1 led to a modest increase in visible pigmentation after 2 hours and a much more dramatic increase after 16 hours compared to the untreated control cells. Similar increases were observed using the less specific deacidification agents ammonium chloride, monensin, or nigericin (Figure S6B). Despite the relative specificity of MME in neutralising lysosomes, the effects of treatment of MME on visible melanin were much more subtle and only detectable when visualised at higher magnification (Figure S6A). This may reflect either a more modest effect of MME on the pH of lysosomes compared to lysosomotropic compounds^72^ or the majority of melanogenesis occurring within esterase-negative late endosomes.

**FIGURE 7.**
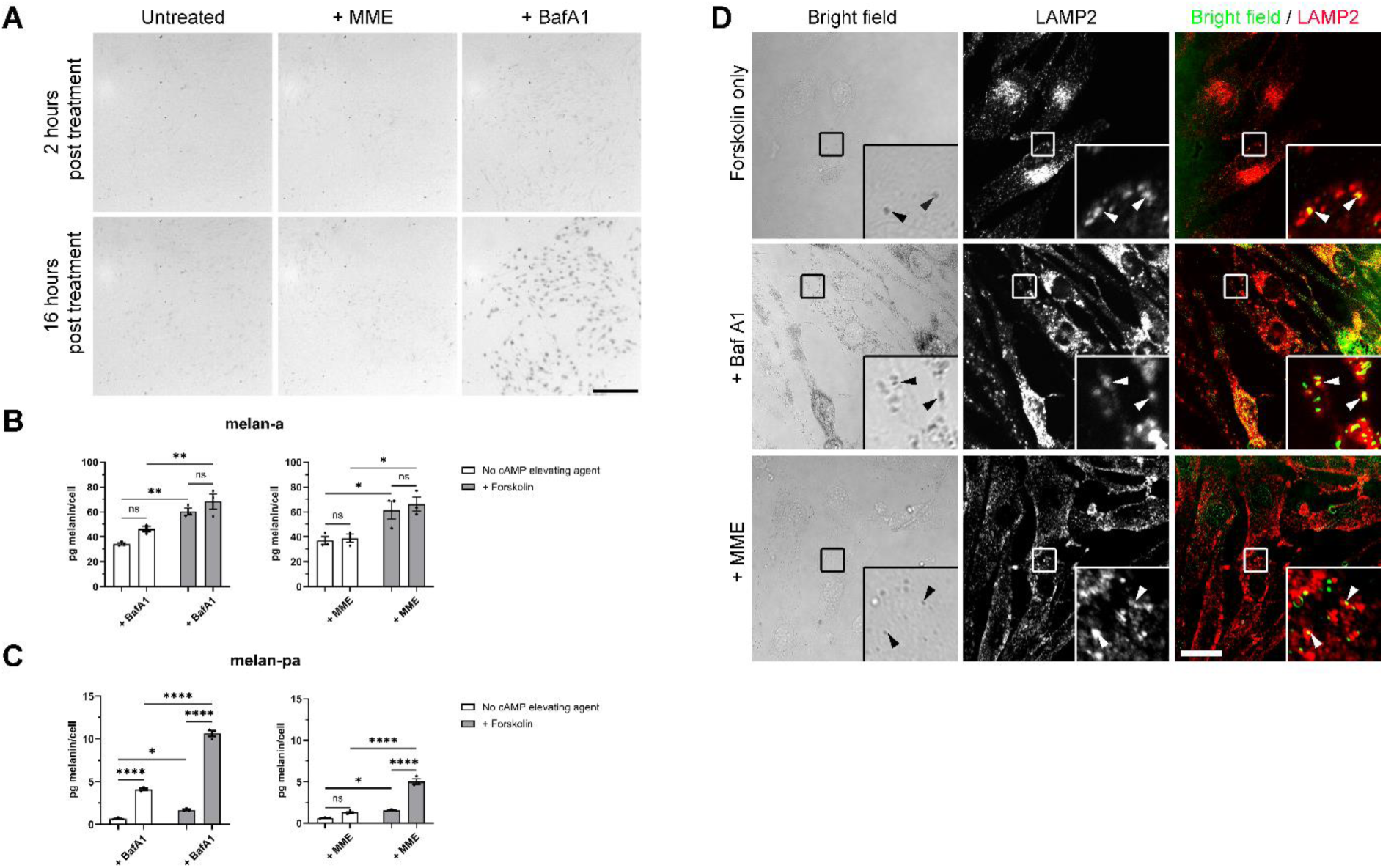
Direct pH modulation by deacidification agents with or without forskolin significantly enhances melanin content in LAMP2-containing compartments of BLOC-1-deficient melanocytes. **A.** Bright field microscopy images of BLOC-1-deficient melanocytes that were untreated or treated with 10 mM MME or 20 nM BafA1 taken at 2 and 16 hours post treatment. Scale bar, 200 µm. **B, C.** Quantitative melanin content analysis of wild-type melan-a melanocytes **(B)** or BLOC-1-deficient melan-pa melanocytes **(C)** cultured for 7 days in the absence or presence of 20 µM forskolin and either 20 nM BafA1 (for final 6 hours) or 10 mM MME (for final 24 hours). Data from 3 independent experiments are expressed as mean pg melanin/cell ± SEM. Statistical significance was determined by two-way ANOVA with a post-hoc Tukey test. Ns, not significant; *, p<0.05; **, p <0.01: ****, p < 0.0001. **D.** BLOC-1-deficient melan-pa melanocytes treated with forskolin for 7 days and either BafA1 (for final 6 hours) or MME for final 24 hours were fixed, immunolabelled for LAMP2 (red), and analysed by confocal IFM (for LAMP2) and bright field microscopy (for pigment granules). Image shows each channel and a merged image in which the bright-field images were inverted and pseudo-coloured green. Insets, boxed regions enlarged 4.5x. Arrowheads show puncta of LAMP2 overlapping with melanin. Scale bar, 20 µm.

We next quantified the effects of BafA1 and MME on melanin content and asked whether the cAMP elevating agent forskolin enhanced these effects. In wild type (Figure 7B) and BLOC-1^R^ melanocytes (melan-mu:MuHA; Figure S6C) neither BafA1 nor MME treatment significantly enhanced basal or forskolin-stimulated melanin content. In contrast, co-treatment of BLOC-1-deficient melan-pa (Figure 7C) or melan-mu (Figure S6D) melanocytes with forskolin and either BafA1 or MME substantially increased melanin content relative to either treatment alone. Taken together these data suggest that direct pH modulation rapidly activates pre-existing tyrosinase and enhance melanin content specifically in BLOC-1-deficient melanocytes.

Lastly, we used IFM and bright field microscopy to test for the localisation of melanin within BLOC-1-deficient melanocytes treated with forskolin and either BafA1 or MME (Figure 7D). While only cells treated with both forskolin and BafA1 harboured abundant well-pigmented melanin containing granules, the melanin in all conditions resided within LAMP-2-containing structures, suggesting that they form in late endosomes/lysosomes and not in melanosomes.

## Discussion

Previous studies in BLOC-1-deficient melanocytes had described melanin in melanosomes with abnormal morphology; small, circular organelles without visible PMEL fibrils but often containing melanin-positive ILVs^47,48,73,74^. Here, we provide evidence to suggest that these organelles are in fact late endosomes or lysosomes. Analysis of the intra-organellar pH (pH_i_) revealed a subpopulation of late endosomes/lysosomes with a pH_i_ that fell within the permissive range for tyrosinase activity (> pH 5.5). Moreover, cargo upregulation due to enhanced MC1R signalling or raised pH_i_ due to genetic or pharmacological treatments led to enhanced melanin deposition in these organelles. Our data support the conclusion that mistargeting of melanosomal cargoes to late endosomes and lysosomes can reprogram these organelles to synthesise melanin under stimulated conditions.

The use of cAMP elevating agents to increase melanin content has previously been established in cell lines and murine models with competent trafficking machineries^20,21,49^. While our present study supports these observations in wild-type and BLOC-1^R^ melanocytes, we also show that BLOC-1-deficient melanocytes retain melanogenic capacity that can be enhanced with cAMP elevating agents despite significant mislocalisation of several melanogenic cargoes. The retention of melanogenic capabilities described here is consistent with both the coat colour of BLOC-1-deficient mice and of skin samples from HPS-mutant mice^47,48,75^, which are not completely devoid of melanin. This contrasts with melanocytes derived from mice with pathogenic tyrosinase mutations that result in the complete absence of melanin^76^. Although the cAMP elevating agent induced increases in the melanin content of BLOC-1-deficient melanocytes are unlikely to be physiologically relevant, they do provide a potential therapeutic avenue for HPS patients and perhaps other trafficking disorders that warrants further research.

In the current study we show that melanin in BLOC-1-deficient melanocytes is predominantly formed not within melanosomes but rather within late endosomes or lysosomes. Melanogenesis within endolysosomes is not an entirely new concept as tyrosinase expressed exogenously within fibroblasts localises to late endosomes and lysosomes and generates melanin within these organelles^77–79^. However, in melanocytes, the melanosomes within which melanin is normally synthesised are distinct from endolysosomal compartments^6^. Nevertheless, previous studies have alluded to melanogenesis in endolysosomal organelles in BLOC-1-deficient melanocytes^47,48,73,74^. These studies show that in BLOC-1-deficient melanocytes melanin resides in small, circular organelles without visible PMEL fibrils but that often contain ILVs filled with melanin. Although these studies generally refer to these organelles as melanosomes, the current study strongly supports the idea that melanogenesis occurs in late endosomes in BLOC-1-deficient melanocytes as melanin resides in LAMP2-positive compartments which, by conventional electron microscopy and CLEM, resemble late endosomal MVBs in morphology; these organelles are smaller and more circular than traditional ellipsoid bona-fide melanosomes. In addition, we noted that ellipsoid striated stage II melanosomes were present in BLOC-1-deficient melanocytes but remain unpigmented, in agreement with previous studies^34,42^. Surprisingly, these compartments largely remained unpigmented even upon treatment with deacidification agents, consistent with a previous study suggesting that these compartments lack the requisite copper cofactor for tyrosinase due to the loss of ATP7A delivery to melanosomes^9^. It is intriguing that the ultrastructure of the MVB-like melanin-containing compartments resembles that of the compartments in which pheomelanin accumulates in melanocytes from red-haired individuals or MC1R mutant animals^80–83^.

How does melanogenesis occur in late endosomes and how do cAMP elevating agents increase the melanin content of BLOC-1-deficient melanocytes? While cAMP elevating agents had little effect on the protein content of tyrosinase and other melanogenic proteins in BLOC-1-competent melanocytes at the time points analysed here, these agents substantially increased melanogenic protein content in BLOC-1-deficient melanocytes, particularly for tyrosinase. Although tyrosinase is crucial for melanogenesis, previous studies have shown that the activity and not the absolute expression level of tyrosinase best correlates with melanin content^13,84–86^. This suggests that the cAMP elevating agent induced increases in melanogenesis in both BLOC-1 competent and BLOC-1-deficient melanocytes may best reflect both increased tyrosinase content and enhanced activity of the tyrosinase present.

Tyrosinase activity requires near neutral pH, and the expression of regulators of melanosome pH, such as OCA2 and SLC45A2, increase in response to cAMP elevating agents^49,66,87^. It is therefore likely that the cAMP-induced increases in melanin of BLOC-1-deficient melanocytes are caused by a combination of increased tyrosinase expression and increased expression of OCA2, SLC45A2, and other proteins that facilitate enzymatic activity, all of which are mislocalised partially to late endosomes in the absence of BLOC-1. We speculate that only a subset of late endosomes accumulates sufficient levels of pH regulators and tyrosinase to both deacidify the normally acidic late endosomes and facilitate tyrosinase activity prior to degradation. This is supported by our data showing that the luminal pH of a cohort of late endosomes/lysosomes is more neutral than the majority of such organelles. Indeed, other cell types harbour lysosomes with heterogeneous luminal pH^60,61^. We show that that the cohort of late endosomes/lysosomes with a more neutral pH can be increased upon treatment with forskolin, likely reflecting enhanced expression of proteins that regulate pH.

We show that partial neutralisation of organelle pH plays a key role in enhancing melanogenesis in BLOC-1-deficient melanocytes. Upon overexpression of OCA2 in BLOC-1-deficient melanocytes a significant increase in melanin content in late endosomes is observed. We speculate that this reflects both the mislocalisation of a cohort of OCA2 to late endosomes and the pH neutralising effect of OCA2 chloride channel activity^10^. In contrast to this the overexpression of SLC45A2 did not significantly enhance melanin content. This may reflect a putative role for SLC45A2 in maintaining the organelle pH gradient that is first established by OCA2^11^ rather than in establishing such a gradient. Direct modulation of organelle pH by treatment with BafA1 or MME rapidly and significantly increased melanin content in BLOC-1-deficient but not wild-type melanocytes. This suggests that melanosome pH is already optimal for melanogenesis within wild-type melanocytes, whereas most late endosomes in BLOC-1-deficient melanocytes are too acidic to support melanogenesis. While the import of copper into melanosomes by the copper transporter ATP7A is necessary for tyrosinase activity in wild-type melanocytes and is disrupted in BLOC-1-deficient melanocytes, excess copper was unable to restore tyrosinase activity at an acidic pH^9^, indicating that pH may also play an important role in the hypopigmentation phenotype of BLOC-1-deficient melanocytes. The current study supports this interpretation, and the cohort of ATP7A that mislocalises to multivesicular late endosomes in these cells^9^ likely supplies sufficient copper cofactor to activate tyrosinase within the deacidified organelles.

In summary we propose that a cohort of inactive tyrosinase is mislocalised to late endosomal MVBs in BLOC-1-deficient melanocytes. Within these organelles, tyrosinase encounters mislocalised BLOC-1-dependent cargoes, including OCA2, ATP7A, and perhaps SLC45A2, that could facilitate melanogenesis within late-endocytic organelles. Increased expression of melanogenic proteins (via enhanced MC1R signalling) and/or modulation of pH (either via enhanced MC1R signalling, genetic manipulation, or the use of deacidification agents) further increases melanin content by facilitating tyrosinase activity in late endosomal MVBs. This work extends our understanding of the hypopigmentation phenotype of BLOC-1-deficient melanocytes and documents a novel mechanism of organelle reprogramming to allow for melanogenesis in the absence of functional melanosomes.

## Acknowledgements

The authors are grateful to Dr Matthew Hannah and Dr Heather Brooks for practical help and advice regarding electron microscopy studies, and to Dr Nyamekye Obeng-Adjei for preliminary studies that led to this work. We are grateful for funding from the Wellcome Trust (grant number 108429/Z/15/Z to E.V.S.) and US National Institutes of Health grant R01EY01625 to M.S.M. (from the National Eye Institute).

## Author Contributions

Conceptualisation, P.S.G., E.V.S., and M.S.M.; Methodology, P.S.G., T.C., M.S.M., and E.V.S.; Investigation, P.S.G and S.P.; Resources T.C. and M.S.M.; Writing – Original Draft, P.S.G.; Writing – Review & Editing, P.S.G., S.P., T.C., M.S.M., and E.V.S.; Visualisation, P.S.G.; Supervision, T.C., E.V.S., and M.S.M.; Funding Acquisition, E.V.S. and M.S.M.

## Materials and Methods

### Cell culture

BLOC-1-deficient murine melanocytes, melan-pa and melan-mu^34^, wild type melan-a^88^, BLOC-1-rescued (melan-mu:MuHA)^34^ and the albino line melan-c2 (unpublished) were maintained in RPMI 1640 supplemented with 2 mM L-glutamine, 10,000 units/ml penicillin, 100 μg/ml streptomycin sulphate and 10% fetal bovine serum. Cells were maintained in this growth medium along with 200 nM 12-0-tetradecanoyl phorbol acetate (TPA) and either 200 pM cholera toxin (CT), 100 pM 4-Norleucine,7-D-phenylalanine-alpha-melanocyte-stimulating hormone (NDP-MSH), or 20 μM forskolin for 7 days unless otherwise stated. For treatment with deacidification agents, BLOC-1-deficient melanocytes were cultured with or without 20 μM forskolin for 6 days, before the medium was changed to contain 20 mM HEPES with or without 20 μM forskolin, 10 mM MME, and/or 20 mM BafA1. Based on preliminary survival curves, cells were treated for 24 hours with MME or 6 hours with BafA1 before harvesting samples to quantify the melanin content as described.

### Antibodies

For IFM studies primary antibodies used were TA99/Mel-5 against TYRP1 (ATCC)^89^, anti-HA tag (ab9110, Abcam) and GL2A7 against LAMP2 (Developmental Studies Hybridoma Bank, USA). Secondary antibodies used were either AlexaFluor488 goat anti-mouse (Life Technologies), AlexaFluor568 goat anti-rabbit (Life Technologies) or AlexaFluor568 goat anti-rat (Life Technologies). For western blotting, we used primary antibodies pep7h (against the c-terminus of tyrosinase;^90^), pep13h (against the c-terminus of Pmel17;^91^), TYRP1 (G-9) (Santa Cruz Biotechnology (sc-166857)), and anti-actin (20-33) (Sigma-Aldrich (A5060)). Secondary antibodies used were IRDye 800CW Goat anti-Mouse IgG (Li-Cor (926-32210)) and IR Dye 680RD Goat anti-Rabbit IgG (Li-Cor (926-68071).

### Intracellular melanin content

Murine melanocytes were plated at 1 x 10^4^ cells/ml into 35 mm (pigmented melanocytes) or 10 cm dishes (BLOC-1-deficient/albino melanocytes) and treated as described above under Cell Culture. Melanin content was quantified based on a method previously described^92^. Briefly, cells were harvested and resuspended in culture medium with an aliquot taken for determination of cell number. The remaining cells were centrifuged at 16,200 x g for 15 minutes at 4°C with the supernatant removed. The cells were lysed with 1% v/v Triton X-100 in complete phosphate-buffered saline lacking calcium and magnesium (cPBS) for 30 minutes at 4°C. The lysate was centrifuged at 16,200 x g for 30 minutes at 4°C. The insoluble fraction containing the melanin was resuspended in 1M NaOH/10% v/v DMSO and incubated at 85°C for at least 30 minutes or until the melanin was fully dissolved. The solubilised melanin was read against a blank of 1M NaOH/10 % v/v DMSO at 405 nm in a SpectraMax 340 spectrophotometer (Molecular Devices). The absorbance was compared to a standard curve of absorbance of known concentrations of synthetic melanin to determine melanin content. The melanin concentration was normalised to the total cell number of each replicate with the background absorbance values of the unpigmented melanocyte line, melan-c2, subtracted (see Figure S1A).

### Western blotting and quantification

Cells were cultured in 10 cm dishes with or without cAMP elevating agents as described above. After 7 days in culture, medium was removed, dishes were placed on ice and washed with ice-cold phosphate-buffered saline without Ca or Mg (PBSA). Cells were lysed with 500µl of ice-cold lysis buffer (50 mM Tris pH 8.0, 150 mM NaCl, 2 mM EDTA, 1% v/v Triton-X 100, 1% w/v sodium dodecyl sulphate (SDS)) containing complete protease inhibitor (Roche), and the lysate was homogenised using a 21-gauge needle and clarified by centrifugation at 16,200 x g for 15 minutes at 4°C. Total protein concentration was determined before separation by SDS-PAGE and transfer to a polyvinylidene difluoride (PVDF)-FL membrane. The membrane was incubated with primary antibodies (see above) overnight diluted in 5% non-fat milk in TBS-T (20 mM Tris, 137 mM sodium chloride, 0.1% v/v Tween-20, pH 7.6) followed by incubation with fluorescently-labelled secondary antibodies (see above) diluted in 5% non-fat milk in TBS-T before detection using the LiCor Odyssey Clx (LI-COR Biosciences). To quantify changes in protein expression between samples, densitometric analysis was performed using Image Studio (Version 5.2, Li-Cor), The fluorescent intensities of each band were measured and the background values subtracted. Relative protein expression was determined using a method previously described^93^.

### Immunolabelling and Immunofluorescence Microscopy

Melanocytes were plated onto glass coverslips at 3 x 10^4^ cells/ml and treated with cAMP elevating agents as described above or transfected as described below. Cells were fixed for 20 minutes with 4% paraformaldehyde in cPBS, rinsed with cPBS and incubated for 5 minutes with 50 mM ammonium chloride. Permeabilisation/blocking buffer (PSS:0.05% v/v saponin, 2% FBS v/v in cPBS) was added to the coverslips for 5 minutes at room temperature and then cells were labelled with primary antibodies (see above) prepared in PSS for 1 hour before washing and subsequent incubation with fluorescently-conjugated secondary antibodies for 1 hour. The cells were washed 3 times with PSS and sometimes incubated with 1 µg/ml DAPI in the dark for 10 minutes. The cells were then washed 3 times with cPBS, twice with water before mounting onto microscope slides with CitiFluor AF1 mounting solution (Agar Scientific). The cells were imaged using a Nikon A1R inverted confocal microscope with NIS Elements software (Version 4.2; Nikon). Images were prepared for presentation using Adobe Photoshop CS6 Version 13.0 (Adobe Systems).

### Conventional electron microscopy and quantification of pigmented organelles

BLOC-1-deficient melan-mu and BLOC-1^R^ melan-mu:MuHA melanocytes were plated at 3 x 10^4^ cells/ml in 10 cm dishes and cultured with TPA in the presence or absence of forskolin as described above. The medium was removed, cells were washed in cPBS and fixed with 0.5% Karnovsky’s fixative (4% w/v PFA, 72 mM sodium cacodylate, 4 mM calcium chloride, 0.5% w/v glutaraldehyde) for 1-2 hours. This was removed, replaced with 2% Karnovsky’s fixative (4% w/v PFA, 72 mM sodium cacodylate, 4 mM calcium chloride, 2% w/v glutaraldehyde) and left overnight. The cells were scraped, centrifuged at 96 x g for 5 minutes, washed twice with 0.1 M sodium cacodylate for 15 minutes, and incubated with 2% w/v osmium tetroxide for 1 hour. Cells were washed twice for 15 minutes each with 0.1 M sodium cacodylate, dehydrated using an ethanol series (2 x 15 minutes in 70% ethanol, 2 x 15 minutes in 90% ethanol and 3 x 30 minutes in absolute ethanol) and then washed 3 times with propylene oxide for 10 minutes each. The samples were infiltrated with a 1:1 mix of propylene oxide and resin (consisting of 36 g Agar 100 (Agar Scientific), 24 g dodecenylsuccinic anhydride (DDSA, Agar Scientific), 15 g methyl nadic anhydride (MNA, Agar Scientific) and 1.13 g of benzyl dimethylamine (BDMA, Agar Scientific)) for 1 hour. This was replaced with 100% resin and left for a further hour. The resin was replaced, and pellets were left to polymerise at 60°C for 48 hours.

Once polymerised, the block was trimmed and ultra-thin sections (approx. 90 nm) were taken using a UltraCutE ultramicrotome (Reichert/Leica) and transferred to copper mesh grids (Agar Scientific). The ultra-thin sections were post-stained by inverting the grids onto a drop of 2% v/v uranyl acetate for 10 minutes at room temperature in the dark followed by 5 30 second washes in dH2O. The grid was then placed onto a drop of Reynolds’ lead citrate^94^ for 90 seconds, washed 5 times for 30 seconds each in dH2O and left to dry before imaging. Images were taken on a Hitachi TEM H-7100 with an AMT XR-16 camera using AMT Capture Engine software (Version V602.580.20d). The maximum length and width of individual melanosomes/melanin-containing organelles was measured from electron microscopy images using the AMT Capture Engine software (Version V602.580.20d). A minimum of 100 such organelles were counted for either melan-mu:MuHA (untreated or treated with forskolin) and melan-mu (untreated or treated with forskolin) from a minimum of 4 preparations each.

### Correlative light and electron microscopy (CLEM)

BLOC-1-deficient melan-mu were plated at 1x10^4^ cells/ml onto 35 mm dishes which have a 1.5 gridded coverslip attached with a raised alphanumeric pattern (MatTek Corporation, USA). Cells were cultured in the presence or absence of forskolin as described above for 7 days. Cells were then washed twice with cPBS prior to fixation with 4% w/v PFA and 0.025% v/v glutaraldehyde for 15 minutes at room temperature. The fixative was removed, and cells were immunolabelled for fluorescence microscopy using LAMP2 as described above, except that after incubation with the secondary antibody, the dishes were incubated with 1 μg/ml DAPI for 10 minutes in the dark and washed 3 times with cPBS for 5 minutes each. Dishes were left in cPBS for imaging.

Dishes were imaged on a Nikon A1R inverted confocal microscope with the NIS Elements software (Version 4.2). Regions of interest were imaged at 60 x magnification, noting the alphanumeric coordinates of these regions to locate the same region once cells had been embedded for electron microscopy. After confocal imaging, the cells were further fixed using Karnovsky’s fixative as described above. Cells were then washed twice in 0.1 M sodium cacodylate for 5 minutes each and incubated with 1.5% w/v potassium ferricyanide and 1% osmium tetroxide (Agar Scientific) for 1 hour on ice. This was followed by 3 x 5-minute washes with 0.1 M sodium cacodylate and subsequent incubation with 1% w/v tannic acid (TAAB Laboratories Equipment Limited) for 45 minutes at room temperature. The cells were washed and incubated with 1% w/v sodium sulphate for 5 minutes prior to dehydration in an ethanol series as described above.

Following dehydration cells were infiltrated with a 1:1 mix of absolute ethanol and resin (composed of 12 g TAAB 812 resin (TAAB Laboratories Equipment Limited), 4.75 g DDSA (TAAB Laboratories Equipment Limited), 8.25 g MNA (TAAB Laboratories Equipment Limited) and 0.75 g of BDMA (TAAB Laboratories Equipment Limited)) for 1 hour. The resin was replaced with 100% resin for 2 hours before the resin was replaced and left for another 2 hours. The gridded coverslips attached to the dish was placed onto the bottom of a sawn-off 15ml tube containing resin before polymerisation at 60°C for 48 hours.

Once polymerised the dish was removed from the tube leaving the cells embedded in the resin along with the alphanumeric coordinates. The coordinates of regions of interest were identified on the polymerised resin surface prior to sectioning by Dr Heather Brooks (Image Resource Facility, St. George’s, University of London). All sections were collected and placed on Formvar coated grids and post-stained as described above. Images were taken on a Hitachi TEM H-7100 with an AMT XR-16 camera using AMT Capture Engine software (Version V602.580.20d). Low magnification images were taken to identify the cell(s) of interest on the section and the orientation relative to the grid. Higher magnification images were taken for presentation. To correlate the light and electron microscopy images, an electron micrograph of an area of interest was chosen and the corresponding area within the light microscopy image was enlarged using Adobe Photoshop CS6 (Version 13.0), so the area/cell(s) were the same size as those in the electron microscopy image. These images were then correlated by identifying specific features (e.g., membrane protrusions, nucleus etc.) within both images and aligning them accordingly using Adobe Photoshop CS6. The light microscopy image was overlaid onto the electron micrograph using the ‘Screen’ function in Adobe Photoshop.

### Transfection and quantification

BLOC-1-deficient melanocytes were cultured on glass coverslips without cAMP-elevating agents as described above. When cells were ∼70% confluent they were transfected with 500 ng of either OCA2 (pCR3/OCA2-HA^63^) or SLC45A2 (pCI-HA-SLC45A2^11^) using Lipofectamine 3000 as per the manufacturer’s instructions. After 48 hours the cells were fixed and processed for IFM as described above. The identity of the cells and plasmids used on the coverslips was blinded before immunolabelling with anti-LAMP2, anti-HA tag and DAPI and confocal microscopy analysis as described above.. In total at least 150 HA-tag positive cells were imaged for each transgene from at least 3 independent experiments and the percentage of HA-positive cells which were pigmented was quantified.

### Lysosomal pH measurements

*In vitro* calibration curves were generated using the pH sensitive fluorophore Oregon Green conjugated to 10,000 MW dextran as previously described^58,59^. Briefly calibration buffers containing 120 mM KCl, 20 mM NaCl, 1 mM MgSO_4_, 20 mM 4-(2-hydroxyethyl)-1-piperazineethanesulfonic acid (HEPES), 0.5 mM CaCl_2_, and 10 mM glucose were adjusted to different pH values (pH 2 - 8). 50 μg/ml Oregon-Green Dextran was then diluted in each of these different pH buffers, transferred to a black-wall 96 well plate (Greiner Bio-One) and the fluorescent-intensity was measured on a BioTek Synergy LX multi-mode reader at room temperature. Calibration curves were generated by normalising the fluorescent intensity of each pH value to the maximal fluorescent measured at pH 7 and curves were fitted using a four-parameter logistic model^95^ using GraphPad Prism (Version 10.2.1). From this, the pKa and the Hill coefficient were calculated.

*In vivo* calibration curves were generated by culturing melan-a in black-wall 96 well plates (Greiner Bio-One) for 6 days. The medium was removed, and cells were washed once with cPBS and incubated overnight with 250 μg/ml Oregon-Green Dextran in full growth medium. The next day the medium was removed and the cells were washed 3 times with cPBS and incubated with full growth medium for 2 hours to enable the fluorescently-labelled dextrans to reach terminal lysosomes. The medium was then removed, cells were washed twice with cPBS and then were incubated with calibration buffer at pH 7 (as described above). Baseline fluorescent-intensity measurements were recorded using BioTek Synergy LX multi-mode reader at room temperature. The pH 7 calibration buffer was removed and replaced with calibration buffers at various pH values (pH 3.5 - 8.0; as described above) containing 5 μM nigericin and 5 μM monensin as previously described^58^ Fluorescent intensity was measured on a BioTek Synergy LX multi-mode reader at room temperature over 20 minutes to allow for equilibration. Calibration curves were generated as described above.

To measure the pH of lysosomes, melanocytes were first plated at 3 x 10^4^ cells/ml in black-wall 96 well plates (Greiner Bio-One) in the presence or absence of CT, NDP-MSH or forskolin for 6 days. The medium was removed and cells were washed once with cPBS and incubated overnight with 250 μg/ml Oregon-Green Dextran in full growth medium and the relevant cAMP-elevating agent. The next day the medium was removed and the cells were washed 3 times with cPBS and incubated with full growth medium for 2 hours to enable the fluorescently-labelled dextrans to reach terminal lysosomes. The medium was then removed, cells were washed twice with cPBS and then incubated with pH 7 calibration buffer (as described above). Baseline fluorescent-intensity measurements were recorded using BioTek Synergy LX multi-mode reader at room temperature (5 reads with a 30 second interval between each) and then 2 M ammonium chloride was added to determine Fmax. The background fluorescent values of unlabelled cells in pH 7 calibration buffer were subtracted from both the steady-state and Fmax values and lysosomal pH was calculated using an equation previously described^58^:

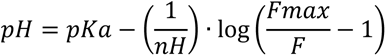

Where *pKA* is pH corresponding to 50% fluorescent intensity, *nH* is the Hill coefficient (both derived from the in vivo calibration curve), *Fmax* is the maximum fluorescent intensity (minus background fluorescence) upon the addition of ammonium chloride and *F* is the steady state fluorescent intensity (minus background fluorescence).

To measure individual pH of lysosomes, BLOC-1-deficient melan-mu melanocytes were plated at 3 x 10^4^ cells/ml in 35 mm glass bottom dishes (VWR) in the presence or absence of 20 µM forskolin for 6 days. The cells were allowed to internalise Oregon-Green Dextran before a 2 hour chase as described above. The medium was then removed, cells were washed twice with cPBS and then incubated in pH 7 calibration buffer (without monensin or nigericin). Images were taken on an Olympus IX71 inverted fluorescence microscope using a U-Plan-Apo 100x 1.35 NA objective with a Ixon3 EMCCD camera (Andor). The EMCCD camera was operated in frame transfer mode at full gain and cooled to -70°C. 512 by 512 pixel images were acquired at 5 frames per second throughout the experiment. Epifluorescence measurements of Oregon-Green Dextran labelled lysosomes were acquired at steady-state and then immediately after exposure to 2 M ammonium chloride (to determine Fmax). The pH of individual lysosomes was derived from these epifluorescence measurements using a previously described macro in ImageJ ^96^ and the parameters describing the relationship between Oregon-Green fluorescence and pH from the calibration curves is shown in Figure S4A. The pH of late endosomes/lysosomes was quantified by measuring the mean intensity of individual organelles from images generated in ImageJ as described above.

## Statistical analysis

Statistical data are presented as mean ± SEM. Statistics were calculated in GraphPad Prism (version 10.2.1) using one-way ANOVA with Dunnett’s multiple comparison test, two-tailed student’s t-test or two-way ANOVA with a post-hoc Tukey test as specified in the figure legend. A p value < 0.05 was considered as statistically significant.

## Supplemental Information

Supplemental PDF. Figures S1-S6.

## References

1. Bowman, S.L., Bi-Karchin, J., Le, L., and Marks, M.S. (2019). The road to lysosome-related organelles: Insights from Hermansky-Pudlak syndrome and other rare diseases. Traffic 20, 404–435. 10.1111/tra.12646.

2. Kobayashi, N., Nakagawa, A., Muramatsu, T., Yamashina, Y., Shirai, T., Hashimoto, M.W., Ishigaki, Y., Ohnishi, T., and Mori, T. (1998). Supranuclear melanin caps reduce ultraviolet induced DNA photoproducts in human epidermis. J Invest Dermatol 110, 806–810. 10.1046/j.1523-1747.1998.00178.x.

3. Graham, D.G., Tiffany, S.M., and Vogel, F.S. (1978). The toxicity of melanin precursors. J Invest Dermatol 70, 113–116. 10.1111/1523-1747.ep12541249.

4. Hochstein, P., and Cohen, G. (1963). The Cytotoxicity of Melanin Precursors. Annals of the New York Academy of Sciences 100, 876–886. 10.1111/j.1749-6632.1963.tb42896.x.

5. Seiji, M., Fitzpatrick, T.B., Simpson, R.T., and Birbeck, M.S.C. (1963). Chemical Composition and Terminology Of Specialized Organelles (Melanosomes and Melanin Granules) in Mammalian Melanocytes. Nature 197, 1082–1084. 10.1038/1971082a0.

6. Raposo, G., Tenza, D., Murphy, D.M., Berson, J.F., and Marks, M.S. (2001). Distinct protein sorting and localization to premelanosomes, melanosomes, and lysosomes in pigmented melanocytic cells. J Cell Biol 152, 809–824. 10.1083/jcb.152.4.809.

7. Hurbain, I., Geerts, W.J., Boudier, T., Marco, S., Verkleij, A.J., Marks, M.S., and Raposo, G. (2008). Electron tomography of early melanosomes: implications for melanogenesis and the generation of fibrillar amyloid sheets. Proc Natl Acad Sci U S A 105, 19726–19731. 10.1073/pnas.0803488105.

8. Hellstrom, A.R., Watt, B., Fard, S.S., Tenza, D., Mannstrom, P., Narfstrom, K., Ekesten, B., Ito, S., Wakamatsu, K., Larsson, J., et al. (2011). Inactivation of Pmel alters melanosome shape but has only a subtle effect on visible pigmentation. PLoS Genet 7, e1002285. 10.1371/journal.pgen.1002285.

9. Setty, S.R., Tenza, D., Sviderskaya, E.V., Bennett, D.C., Raposo, G., and Marks, M.S. (2008). Cell-specific ATP7A transport sustains copper-dependent tyrosinase activity in melanosomes. Nature 454, 1142–1146. 10.1038/nature07163.

10. Bellono, N.W., Escobar, I.E., Lefkovith, A.J., Marks, M.S., and Oancea, E. (2014). An intracellular anion channel critical for pigmentation. Elife 3, e04543. 10.7554/eLife.04543.

11. Le, L., Escobar, I.E., Ho, T., Lefkovith, A.J., Latteri, E., Haltaufderhyde, K.D., Dennis, M.K., Plowright, L., Sviderskaya, E.V., Bennett, D.C., et al. (2020). SLC45A2 protein stability and regulation of melanosome pH determine melanocyte pigmentation. Mol Biol Cell 31, 2687–2702. 10.1091/mbc.E20-03-0200.

12. Ancans, J., Tobin, D.J., Hoogduijn, M.J., Smit, N.P., Wakamatsu, K., and Thody, A.J. (2001). Melanosomal pH controls rate of melanogenesis, eumelanin/phaeomelanin ratio and melanosome maturation in melanocytes and melanoma cells. Exp Cell Res 268, 26–35. 10.1006/excr.2001.5251.

13. Fuller, B.B., Spaulding, D.T., and Smith, D.R. (2001). Regulation of the catalytic activity of preexisting tyrosinase in black and Caucasian human melanocyte cell cultures. Exp Cell Res 262, 197–208. 10.1006/excr.2000.5092.

14. Bertolotto, C., Abbe, P., Hemesath, T.J., Bille, K., Fisher, D.E., Ortonne, J.P., and Ballotti, R. (1998). Microphthalmia gene product as a signal transducer in cAMP-induced differentiation of melanocytes. J Cell Biol 142, 827–835. 10.1083/jcb.142.3.827.

15. Price, E.R., Horstmann, M.A., Wells, A.G., Weilbaecher, K.N., Takemoto, C.M., Landis, M.W., and Fisher, D.E. (1998). alpha-Melanocyte-stimulating hormone signaling regulates expression of microphthalmia, a gene deficient in Waardenburg syndrome. J Biol Chem 273, 33042–33047. 10.1074/jbc.273.49.33042.

16. Bertolotto, C., Busca, R., Abbe, P., Bille, K., Aberdam, E., Ortonne, J.P., and Ballotti, R. (1998). Different cis-acting elements are involved in the regulation of TRP1 and TRP2 promoter activities by cyclic AMP: pivotal role of M boxes (GTCATGTGCT) and of microphthalmia. Mol Cell Biol 18, 694–702. 10.1128/MCB.18.2.694.

17. Levine, N., Sheftel, S.N., Eytan, T., Dorr, R.T., Hadley, M.E., Weinrach, J.C., Ertl, G.A., Toth, K., McGee, D.L., and Hruby, V.J. (1991). Induction of skin tanning by subcutaneous administration of a potent synthetic melanotropin. JAMA 266, 2730–2736.

18. Barnetson, R.S., Ooi, T.K., Zhuang, L., Halliday, G.M., Reid, C.M., Walker, P.C., Humphrey, S.M., and Kleinig, M.J. (2006). [Nle4-D-Phe7]-alpha-melanocyte-stimulating hormone significantly increased pigmentation and decreased UV damage in fair-skinned Caucasian volunteers. J Invest Dermatol 126, 1869–1878. 10.1038/sj.jid.5700317.

19. Bautista, R.M., Carter, K.M., Jarrett, S.G., Napier, D., Wakamatsu, K., Ito, S., and D’Orazio, J.A. (2020). Cutaneous pharmacologic cAMP induction induces melanization of the skin and improves recovery from ultraviolet injury in melanocortin 1 receptor-intact or heterozygous skin. Pigment Cell Melanoma Res 33, 30–40. 10.1111/pcmr.12817.

20. D’Orazio, J.A., Nobuhisa, T., Cui, R., Arya, M., Spry, M., Wakamatsu, K., Igras, V., Kunisada, T., Granter, S.R., Nishimura, E.K., et al. (2006). Topical drug rescue strategy and skin protection based on the role of Mc1r in UV-induced tanning. Nature 443, 340–344. 10.1038/nature05098.

21. Spry, M.L., Vanover, J.C., Scott, T., Abona-Ama, O., Wakamatsu, K., Ito, S., and D’Orazio, J.A. (2009). Prolonged treatment of fair-skinned mice with topical forskolin causes persistent tanning and UV protection. Pigment Cell Melanoma Res 22, 219–229. 10.1111/j.1755-148X.2008.00536.x.

22. Hermansky, F., and Pudlak, P. (1959). Albinism associated with hemorrhagic diathesis and unusual pigmented reticular cells in the bone marrow: report of two cases with histochemical studies. Blood 14, 162–169. 10.1182/blood-2016-01-696435.

23. Ammann, S., Schulz, A., Krageloh-Mann, I., Dieckmann, N.M., Niethammer, K., Fuchs, S., Eckl, K.M., Plank, R., Werner, R., Altmuller, J., et al. (2016). Mutations in AP3D1 associated with immunodeficiency and seizures define a new type of Hermansky-Pudlak syndrome. Blood 127, 997–1006. 10.1182/blood-2015-09-671636.

24. Anikster, Y., Huizing, M., White, J., Shevchenko, Y.O., Fitzpatrick, D.L., Touchman, J.W., Compton, J.G., Bale, S.J., Swank, R.T., Gahl, W.A., and Toro, J.R. (2001). Mutation of a new gene causes a unique form of Hermansky-Pudlak syndrome in a genetic isolate of central Puerto Rico. Nat Genet 28, 376–380. 10.1038/ng576.

25. Badolato, R., Prandini, A., Caracciolo, S., Colombo, F., Tabellini, G., Giacomelli, M., Cantarini, M.E., Pession, A., Bell, C.J., Dinwiddie, D.L., et al. (2012). Exome sequencing reveals a pallidin mutation in a Hermansky-Pudlak-like primary immunodeficiency syndrome. Blood 119, 3185–3187. 10.1182/blood-2012-01-404350.

26. Dell’Angelica, E.C., Shotelersuk, V., Aguilar, R.C., Gahl, W.A., and Bonifacino, J.S. (1999). Altered trafficking of lysosomal proteins in Hermansky-Pudlak syndrome due to mutations in the beta 3A subunit of the AP-3 adaptor. Mol Cell 3, 11–21. 10.1016/s1097-2765(00)80170-7.

27. Li, W., Zhang, Q., Oiso, N., Novak, E.K., Gautam, R., O’Brien, E.P., Tinsley, C.L., Blake, D.J., Spritz, R.A., Copeland, N.G., et al. (2003). Hermansky-Pudlak syndrome type 7 (HPS-7) results from mutant dysbindin, a member of the biogenesis of lysosome-related organelles complex 1 (BLOC-1). Nat Genet 35, 84–89. 10.1038/ng1229.

28. Morgan, N.V., Pasha, S., Johnson, C.A., Ainsworth, J.R., Eady, R.A., Dawood, B., McKeown, C., Trembath, R.C., Wilde, J., Watson, S.P., and Maher, E.R. (2006). A germline mutation in BLOC1S3/reduced pigmentation causes a novel variant of Hermansky-Pudlak syndrome (HPS8). Am J Hum Genet 78, 160–166. 10.1086/499338.

29. Oh, J., Bailin, T., Fukai, K., Feng, G.H., Ho, L., Mao, J.I., Frenk, E., Tamura, N., and Spritz, R.A. (1996). Positional cloning of a gene for Hermansky-Pudlak syndrome, a disorder of cytoplasmic organelles. Nat Genet 14, 300–306. 10.1038/ng1196-300.

30. Pennamen, P., Le, L., Tingaud-Sequeira, A., Fiore, M., Bauters, A., Van Duong Beatrice, N., Coste, V., Bordet, J.C., Plaisant, C., Diallo, M., et al. (2020). BLOC1S5 pathogenic variants cause a new type of Hermansky-Pudlak syndrome. Genet Med 22, 1613–1622. 10.1038/s41436-020-0867-5.

31. Suzuki, T., Li, W., Zhang, Q., Karim, A., Novak, E.K., Sviderskaya, E.V., Hill, S.P., Bennett, D.C., Levin, A.V., Nieuwenhuis, H.K., et al. (2002). Hermansky-Pudlak syndrome is caused by mutations in HPS4, the human homolog of the mouse light-ear gene. Nat Genet 30, 321–324. 10.1038/ng835.

32. Zhang, Q., Zhao, B., Li, W., Oiso, N., Novak, E.K., Rusiniak, M.E., Gautam, R., Chintala, S., O’Brien, E.P., Zhang, Y., et al. (2003). Ru2 and Ru encode mouse orthologs of the genes mutated in human Hermansky-Pudlak syndrome types 5 and 6. Nat Genet 33, 145–153. 10.1038/ng1087.

33. 33. Di Pietro, S.M., Falcon-Perez, J.M., Tenza, D., Setty, S.R., Marks, M.S., Raposo, G., and Dell’Angelica, E.C. (2006). BLOC-1 interacts with BLOC-2 and the AP-3 complex to facilitate protein trafficking on endosomes. Mol Biol Cell 17, 4027–4038. 10.1091/mbc.e06-05-0379.

34. Setty, S.R., Tenza, D., Truschel, S.T., Chou, E., Sviderskaya, E.V., Theos, A.C., Lamoreux, M.L., Di Pietro, S.M., Starcevic, M., Bennett, D.C., et al. (2007). BLOC-1 is required for cargo-specific sorting from vacuolar early endosomes toward lysosome-related organelles. Mol Biol Cell 18, 768–780. 10.1091/mbc.e06-12-1066.

35. Sitaram, A., Dennis, M.K., Chaudhuri, R., De Jesus-Rojas, W., Tenza, D., Setty, S.R., Wood, C.S., Sviderskaya, E.V., Bennett, D.C., Raposo, G., et al. (2012). Differential recognition of a dileucine-based sorting signal by AP-1 and AP-3 reveals a requirement for both BLOC-1 and AP-3 in delivery of OCA2 to melanosomes. Mol Biol Cell 23, 3178–3192. 10.1091/mbc.E11-06-0509.

36. Ciciotte, S.L., Gwynn, B., Moriyama, K., Huizing, M., Gahl, W.A., Bonifacino, J.S., and Peters, L.L. (2003). Cappuccino, a mouse model of Hermansky-Pudlak syndrome, encodes a novel protein that is part of the pallidin-muted complex (BLOC-1). Blood 101, 4402–4407. 10.1182/blood-2003-01-0020.

37. Falcon-Perez, J.M., Starcevic, M., Gautam, R., and Dell’Angelica, E.C. (2002). BLOC-1, a novel complex containing the pallidin and muted proteins involved in the biogenesis of melanosomes and platelet-dense granules. J Biol Chem 277, 28191–28199. 10.1074/jbc.M204011200.

38. Gwynn, B., Martina, J.A., Bonifacino, J.S., Sviderskaya, E.V., Lamoreux, M.L., Bennett, D.C., Moriyama, K., Huizing, M., Helip-Wooley, A., Gahl, W.A., et al. (2004). Reduced pigmentation (rp), a mouse model of Hermansky-Pudlak syndrome, encodes a novel component of the BLOC-1 complex. Blood 104, 3181–3189. 10.1182/blood-2004-04-1538.

39. Moriyama, K., and Bonifacino, J.S. (2002). Pallidin is a component of a multi-protein complex involved in the biogenesis of lysosome-related organelles. Traffic 3, 666–677. 10.1034/j.1600-0854.2002.30908.x.

40. Starcevic, M., and Dell’Angelica, E.C. (2004). Identification of snapin and three novel proteins (BLOS1, BLOS2, and BLOS3/reduced pigmentation) as subunits of biogenesis of lysosome-related organelles complex-1 (BLOC-1). J Biol Chem 279, 28393–28401. 10.1074/jbc.M402513200.

41. Lee, H.H., Nemecek, D., Schindler, C., Smith, W.J., Ghirlando, R., Steven, A.C., Bonifacino, J.S., and Hurley, J.H. (2012). Assembly and architecture of biogenesis of lysosome-related organelles complex-1 (BLOC-1). J Biol Chem 287, 5882–5890. 10.1074/jbc.M111.325746.

42. Delevoye, C., Heiligenstein, X., Ripoll, L., Gilles-Marsens, F., Dennis, M.K., Linares, R.A., Derman, L., Gokhale, A., Morel, E., Faundez, V., et al. (2016). BLOC-1 Brings Together the Actin and Microtubule Cytoskeletons to Generate Recycling Endosomes. Curr Biol 26, 1–13. 10.1016/j.cub.2015.11.020.

43. Delevoye, C., Miserey-Lenkei, S., Montagnac, G., Gilles-Marsens, F., Paul-Gilloteaux, P., Giordano, F., Waharte, F., Marks, M.S., Goud, B., and Raposo, G. (2014). Recycling endosome tubule morphogenesis from sorting endosomes requires the kinesin motor KIF13A. Cell Rep 6, 445–454. 10.1016/j.celrep.2014.01.002.

44. Jani, R.A., Di Cicco, A., Keren-Kaplan, T., Vale-Costa, S., Hamaoui, D., Hurbain, I., Tsai, F.C., Di Marco, M., Mace, A.S., Zhu, Y., et al. (2022). PI4P and BLOC-1 remodel endosomal membranes into tubules. J Cell Biol 221. 10.1083/jcb.202110132.

45. Dennis, M.K., Delevoye, C., Acosta-Ruiz, A., Hurbain, I., Romao, M., Hesketh, G.G., Goff, P.S., Sviderskaya, E.V., Bennett, D.C., Luzio, J.P., et al. (2016). BLOC-1 and BLOC-3 regulate VAMP7 cycling to and from melanosomes via distinct tubular transport carriers. J Cell Biol 214, 293–308. 10.1083/jcb.201605090.

46. Theos, A.C., Tenza, D., Martina, J.A., Hurbain, I., Peden, A.A., Sviderskaya, E.V., Stewart, A., Robinson, M.S., Bennett, D.C., Cutler, D.F., et al. (2005). Functions of adaptor protein (AP)-3 and AP-1 in tyrosinase sorting from endosomes to melanosomes. Mol Biol Cell 16, 5356–5372. 10.1091/mbc.e05-07-0626.

47. Nguyen, T., Novak, E.K., Kermani, M., Fluhr, J., Peters, L.L., Swank, R.T., and Wei, M.L. (2002). Melanosome morphologies in murine models of hermansky-pudlak syndrome reflect blocks in organelle development. J Invest Dermatol 119, 1156–1164. 10.1046/j.1523-1747.2002.19535.x.

48. Nguyen, T., and Wei, M.L. (2004). Characterization of melanosomes in murine Hermansky-Pudlak syndrome: mechanisms of hypopigmentation. J Invest Dermatol 122, 452–460. 10.1046/j.0022-202X.2004.22117.x.

49. Cheli, Y., Luciani, F., Khaled, M., Beuret, L., Bille, K., Gounon, P., Ortonne, J.P., Bertolotto, C., and Ballotti, R. (2009). alphaMSH and Cyclic AMP elevating agents control melanosome pH through a protein kinase A-independent mechanism. J Biol Chem 284, 18699–18706. 10.1074/jbc.M109.005819.

50. Huizing, M., Sarangarajan, R., Strovel, E., Zhao, Y., Gahl, W.A., and Boissy, R.E. (2001). AP-3 mediates tyrosinase but not TRP-1 trafficking in human melanocytes. Mol Biol Cell 12, 2075–2085. 10.1091/mbc.12.7.2075.

51. Chen, J.W., Murphy, T.L., Willingham, M.C., Pastan, I., and August, J.T. (1985). Identification of two lysosomal membrane glycoproteins. J Cell Biol 101, 85–95. 10.1083/jcb.101.1.85.

52. Luzio, J.P., Hackmann, Y., Dieckmann, N.M., and Griffiths, G.M. (2014). The biogenesis of lysosomes and lysosome-related organelles. Cold Spring Harb Perspect Biol 6, a016840. 10.1101/cshperspect.a016840.

53. Shearer, L.J., and Petersen, N.O. (2019). Distribution and Co-localization of endosome markers in cells. Heliyon 5, e02375. 10.1016/j.heliyon.2019.e02375.

54. Bissig, C., Hurbain, I., Raposo, G., and van Niel, G. (2017). PIKfyve activity regulates reformation of terminal storage lysosomes from endolysosomes. Traffic 18, 747–757. 10.1111/tra.12525.

55. Humphries, W.H.t., Szymanski, C.J., and Payne, C.K. (2011). Endo-lysosomal vesicles positive for Rab7 and LAMP1 are terminal vesicles for the transport of dextran. PLoS One 6, e26626. 10.1371/journal.pone.0026626.

56. Seiji, M., Fitzpatrick, T.B., and Birbeck, M.S. (1961). The melanosome: a distinctive subcellular particle of mammalian melanocytes and the site of melanogenesis. J Invest Dermatol 36, 243–252. 10.1038/jid.1961.42.

57. Halaban, R., Patton, R.S., Cheng, E., Svedine, S., Trombetta, E.S., Wahl, M.L., Ariyan, S., and Hebert, D.N. (2002). Abnormal acidification of melanoma cells induces tyrosinase retention in the early secretory pathway. J Biol Chem 277, 14821–14828. 10.1074/jbc.M111497200.

58. Erent, M., Meli, A., Moisoi, N., Babich, V., Hannah, M.J., Skehel, P., Knipe, L., Zupancic, G., Ogden, D., and Carter, T. (2007). Rate, extent and concentration dependence of histamine-evoked Weibel-Palade body exocytosis determined from individual fusion events in human endothelial cells. J Physiol 583, 195–212. 10.1113/jphysiol.2007.132993.

59. Kneen, M., Farinas, J., Li, Y., and Verkman, A.S. (1998). Green fluorescent protein as a noninvasive intracellular pH indicator. Biophys J 74, 1591–1599. 10.1016/S0006-3495(98)77870-1.

60. Johnson, D.E., Ostrowski, P., Jaumouille, V., and Grinstein, S. (2016). The position of lysosomes within the cell determines their luminal pH. J Cell Biol 212, 677–692. 10.1083/jcb.201507112.

61. Bright, N.A., Davis, L.J., and Luzio, J.P. (2016). Endolysosomes Are the Principal Intracellular Sites of Acid Hydrolase Activity. Curr Biol 26, 2233–2245. 10.1016/j.cub.2016.06.046.

62. Bartolke, R., Heinisch, J.J., Wieczorek, H., and Vitavska, O. (2014). Proton-associated sucrose transport of mammalian solute carrier family 45: an analysis in Saccharomyces cerevisiae. Biochem J 464, 193–201. 10.1042/BJ20140572.

63. Sitaram, A., Piccirillo, R., Palmisano, I., Harper, D.C., Dell’Angelica, E.C., Schiaffino, M.V., and Marks, M.S. (2009). Localization to mature melanosomes by virtue of cytoplasmic dileucine motifs is required for human OCA2 function. Mol Biol Cell 20, 1464–1477. 10.1091/mbc.e08-07-0710.

64. Seberg, H.E., Van Otterloo, E., Loftus, S.K., Liu, H., Bonde, G., Sompallae, R., Gildea, D.E., Santana, J.F., Manak, J.R., Pavan, W.J., et al. (2017). TFAP2 paralogs regulate melanocyte differentiation in parallel with MITF. PLoS Genet 13, e1006636. 10.1371/journal.pgen.1006636.

65. Visser, M., Kayser, M., and Palstra, R.J. (2012). HERC2 rs12913832 modulates human pigmentation by attenuating chromatin-loop formation between a long-range enhancer and the OCA2 promoter. Genome Res 22, 446–455. 10.1101/gr.128652.111.

66. Newton, R.A., Cook, A.L., Roberts, D.W., Leonard, J.H., and Sturm, R.A. (2007). Post-transcriptional regulation of melanin biosynthetic enzymes by cAMP and resveratrol in human melanocytes. J Invest Dermatol 127, 2216–2227. 10.1038/sj.jid.5700840.

67. Ancans, J., Hoogduijn, M.J., and Thody, A.J. (2001). Melanosomal pH, pink locus protein and their roles in melanogenesis. J Invest Dermatol 117, 158–159. 10.1046/j.0022-202x.2001.01397.x.

68. Ancans, J., and Thody, A.J. (2000). Activation of melanogenesis by vacuolar type H(+)-ATPase inhibitors in amelanotic, tyrosinase positive human and mouse melanoma cells. FEBS Lett 478, 57–60. 10.1016/s0014-5793(00)01795-6.

69. Chen, K., Minwalla, L., Ni, L., and Orlow, S.J. (2004). Correction of defective early tyrosinase processing by bafilomycin A1 and monensin in pink-eyed dilution melanocytes. Pigment Cell Res 17, 36–42. 10.1046/j.1600-0749.2003.00106.x.

70. Dooley, C.M., Schwarz, H., Mueller, K.P., Mongera, A., Konantz, M., Neuhauss, S.C., Nusslein-Volhard, C., and Geisler, R. (2013). Slc45a2 and V-ATPase are regulators of melanosomal pH homeostasis in zebrafish, providing a mechanism for human pigment evolution and disease. Pigment Cell Melanoma Res 26, 205–217. 10.1111/pcmr.12053.

71. Ni-Komatsu, L., and Orlow, S.J. (2006). Heterologous expression of tyrosinase recapitulates the misprocessing and mistrafficking in oculocutaneous albinism type 2: effects of altering intracellular pH and pink-eyed dilution gene expression. Exp Eye Res 82, 519–528. 10.1016/j.exer.2005.08.013.

72. Decker, R.S., Decker, M.L., Thomas, V., and Fuseler, J.W. (1985). Responses of cultured cardiac myocytes to lysosomotropic compounds and methylated amino acids. J Cell Sci 74, 119–135. 10.1242/jcs.74.1.119.

73. Ito, M., Hashimoto, K., and Organisciak, D.T. (1982). Ultrastructural, histochemical, and biochemical studies of the melanin metabolism in eye and skin of pallid mice. J Invest Dermatol 78, 414–424. 10.1111/1523-1747.ep12507677.

74. Zhang, Q., Li, W., Novak, E.K., Karim, A., Mishra, V.S., Kingsmore, S.F., Roe, B.A., Suzuki, T., and Swank, R.T. (2002). The gene for the muted (mu) mouse, a model for Hermansky-Pudlak syndrome, defines a novel protein which regulates vesicle trafficking. Hum Mol Genet 11, 697–706. 10.1093/hmg/11.6.697.

75. Gautam, R., Novak, E.K., Tan, J., Wakamatsu, K., Ito, S., and Swank, R.T. (2006). Interaction of Hermansky-Pudlak Syndrome genes in the regulation of lysosome-related organelles. Traffic 7, 779–792. 10.1111/j.1600-0854.2006.00431.x.

76. Bennett, D.C., Cooper, P.J., Dexter, T.J., Devlin, L.M., Heasman, J., and Nester, B. (1989). Cloned mouse melanocyte lines carrying the germline mutations albino and brown: complementation in culture. Development 105, 379–385. 10.1242/dev.105.2.379.

77. Bouchard, B., Fuller, B.B., Vijayasaradhi, S., and Houghton, A.N. (1989). Induction of pigmentation in mouse fibroblasts by expression of human tyrosinase cDNA. J Exp Med 169, 2029–2042. 10.1084/jem.169.6.2029.

78. Winder, A.J. (1991). Expression of a mouse tyrosinase cDNA in 3T3 Swiss mouse fibroblasts. Biochem Biophys Res Commun 178, 739–745. 10.1016/0006-291x(91)90170-c.

79. Calvo, P.A., Frank, D.W., Bieler, B.M., Berson, J.F., and Marks, M.S. (1999). A cytoplasmic sequence in human tyrosinase defines a second class of di-leucine-based sorting signals for late endosomal and lysosomal delivery. J Biol Chem 274, 12780–12789. 10.1074/jbc.274.18.12780.

80. Jimbow, K., Ishida, O., Ito, S., Hori, Y., Witkop, C.J., Jr., and King, R.A. (1983). Combined chemical and electron microscopic studies of pheomelanosomes in human red hair. J Invest Dermatol 81, 506–511. 10.1111/1523-1747.ep12522838.

81. Moyer, F.H. (1966). Genetic Variations in the Fine Structure and Ontogeny of Mouse Melanin Granules. American Zoologist 6, 43–66.

82. Liu, Y., Hong, L., Wakamatsu, K., Ito, S., Adhyaru, B., Cheng, C.Y., Bowers, C.R., and Simon, J.D. (2005). Comparison of structural and chemical properties of black and red human hair melanosomes. Photochem Photobiol 81, 135–144. 10.1562/2004-08-03-RA-259.1.

83. Garcia, R.I., and Szabo, G. (1983). Modulation of melanosome ultrastructure in cultured embryonic pigment cells. J Exp Zool 225, 285–291. 10.1002/jez.1402250211.

84. Cook, A.L., Chen, W., Thurber, A.E., Smit, D.J., Smith, A.G., Bladen, T.G., Brown, D.L., Duffy, D.L., Pastorino, L., Bianchi-Scarra, G., et al. (2009). Analysis of cultured human melanocytes based on polymorphisms within the SLC45A2/MATP, SLC24A5/NCKX5, and OCA2/P loci. J Invest Dermatol 129, 392–405. 10.1038/jid.2008.211.

85. Iozumi, K., Hoganson, G.E., Pennella, R., Everett, M.A., and Fuller, B.B. (1993). Role of tyrosinase as the determinant of pigmentation in cultured human melanocytes. J Invest Dermatol 100, 806–811. 10.1111/1523-1747.ep12476630.

86. Halaban, R., Pomerantz, S.H., Marshall, S., Lambert, D.T., and Lerner, A.B. (1983). Regulation of tyrosinase in human melanocytes grown in culture. J Cell Biol 97, 480–488. 10.1083/jcb.97.2.480.

87. Ostojic, J., Yoon, Y.S., Sonntag, T., Nguyen, B., Vaughan, J.M., Shokhirev, M., and Montminy, M. (2021). Transcriptional co-activator regulates melanocyte differentiation and oncogenesis by integrating cAMP and MAPK/ERK pathways. Cell Rep 35, 109136. 10.1016/j.celrep.2021.109136.

88. Bennett, D.C., Cooper, P.J., and Hart, I.R. (1987). A line of non-tumorigenic mouse melanocytes, syngeneic with the B16 melanoma and requiring a tumour promoter for growth. Int J Cancer 39, 414–418. 10.1002/ijc.2910390324.

89. Thomson, T.M., Mattes, M.J., Roux, L., Old, L.J., and Lloyd, K.O. (1985). Pigmentation-associated glycoprotein of human melanomas and melanocytes: definition with a mouse monoclonal antibody. J Invest Dermatol 85, 169–174. 10.1111/1523-1747.ep12276608.

90. Berson, J.F., Frank, D.W., Calvo, P.A., Bieler, B.M., and Marks, M.S. (2000). A common temperature-sensitive allelic form of human tyrosinase is retained in the endoplasmic reticulum at the nonpermissive temperature. J Biol Chem 275, 12281–12289. 10.1074/jbc.275.16.12281.

91. Berson, J.F., Harper, D.C., Tenza, D., Raposo, G., and Marks, M.S. (2001). Pmel17 initiates premelanosome morphogenesis within multivesicular bodies. Mol Biol Cell 12, 3451–3464. 10.1091/mbc.12.11.3451.

92. Oancea, E., Vriens, J., Brauchi, S., Jun, J., Splawski, I., and Clapham, D.E. (2009). TRPM1 forms ion channels associated with melanin content in melanocytes. Sci Signal 2, ra21. 10.1126/scisignal.2000146.

93. Taylor, S.C., Rosselli-Murai, L.K., Crobeddu, B., and Plante, I. (2022). A critical path to producing high quality, reproducible data from quantitative western blot experiments. Sci Rep 12, 17599. 10.1038/s41598-022-22294-x.

94. Reynolds, E.S. (1963). The use of lead citrate at high pH as an electron-opaque stain in electron microscopy. J Cell Biol 17, 208–212. 10.1083/jcb.17.1.208.

95. Tompkins, L.S., Nullmeyer, K.D., Murphy, S.M., Weber, C.S., and Lynch, R.M. (2002). Regulation of secretory granule pH in insulin-secreting cells. Am J Physiol Cell Physiol 283, C429–437. 10.1152/ajpcell.01066.2000.

96. Meli, A., McCormack, A., Conte, I., Chen, Q., Streetley, J., Rose, M.L., Bierings, R., Hannah, M.J., Molloy, J.E., Rosenthal, P.B., and Carter, T. (2023). Altered Storage and Function of von Willebrand Factor in Human Cardiac Microvascular Endothelial Cells Isolated from Recipient Transplant Hearts. Int J Mol Sci 24. 10.3390/ijms24054553.

