## Supplemental Figures for "Enhanced MC1R-signalling and pH modulation facilitate melanogenesis within late endosomes of BLOC-1-deficient melanocytes"

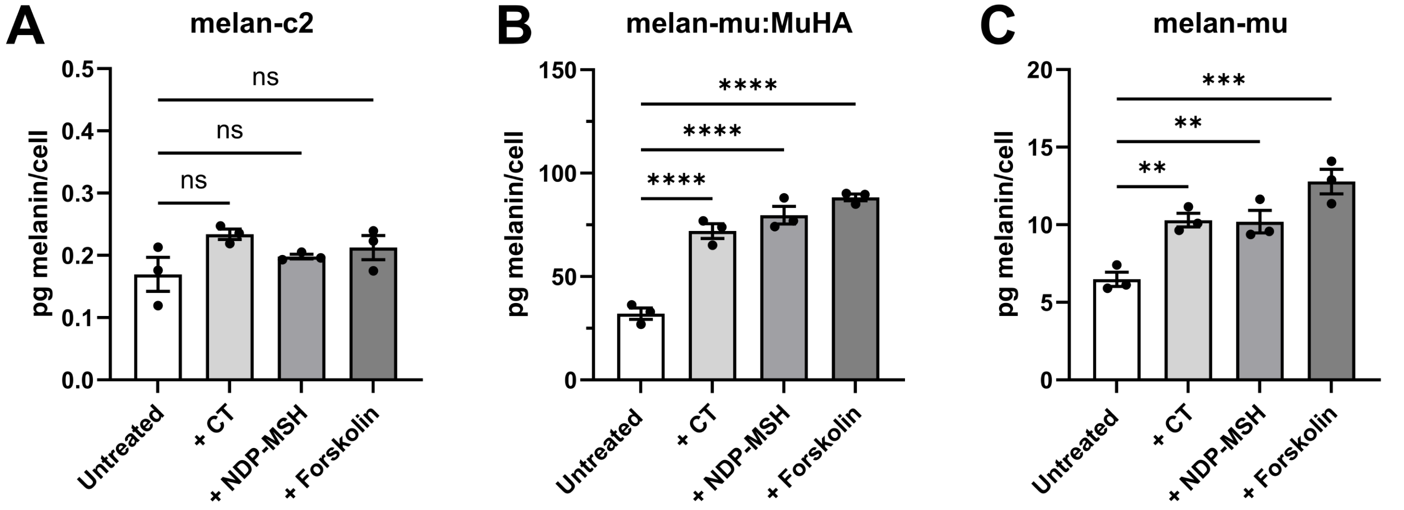


**FIGURE S1. cAMP elevating agents increase melanin content of BLOC-1$^{R}$ and BLOC-1-deficient melanocytes but not tyrosinase-deficient melan-c2**

Quantification of melanin content of melan-c2 **(A)**, melan-mu:MuHA **(B)** and melan-mu **(C)** cultured for 7 days in the presence or absence of either 200 nM CT, 100 pM NDP-MSH or 20 µM forskolin. Data are from 3 independent experiments expressed as mean pg melanin/cell ± SEM. Statistical significance relative to the untreated control was determined by one-way ANOVA with Dunnett's multiple comparison test. Ns, not significant; **, p<0.01; ***, p<0.001; ****, p<0.0001.

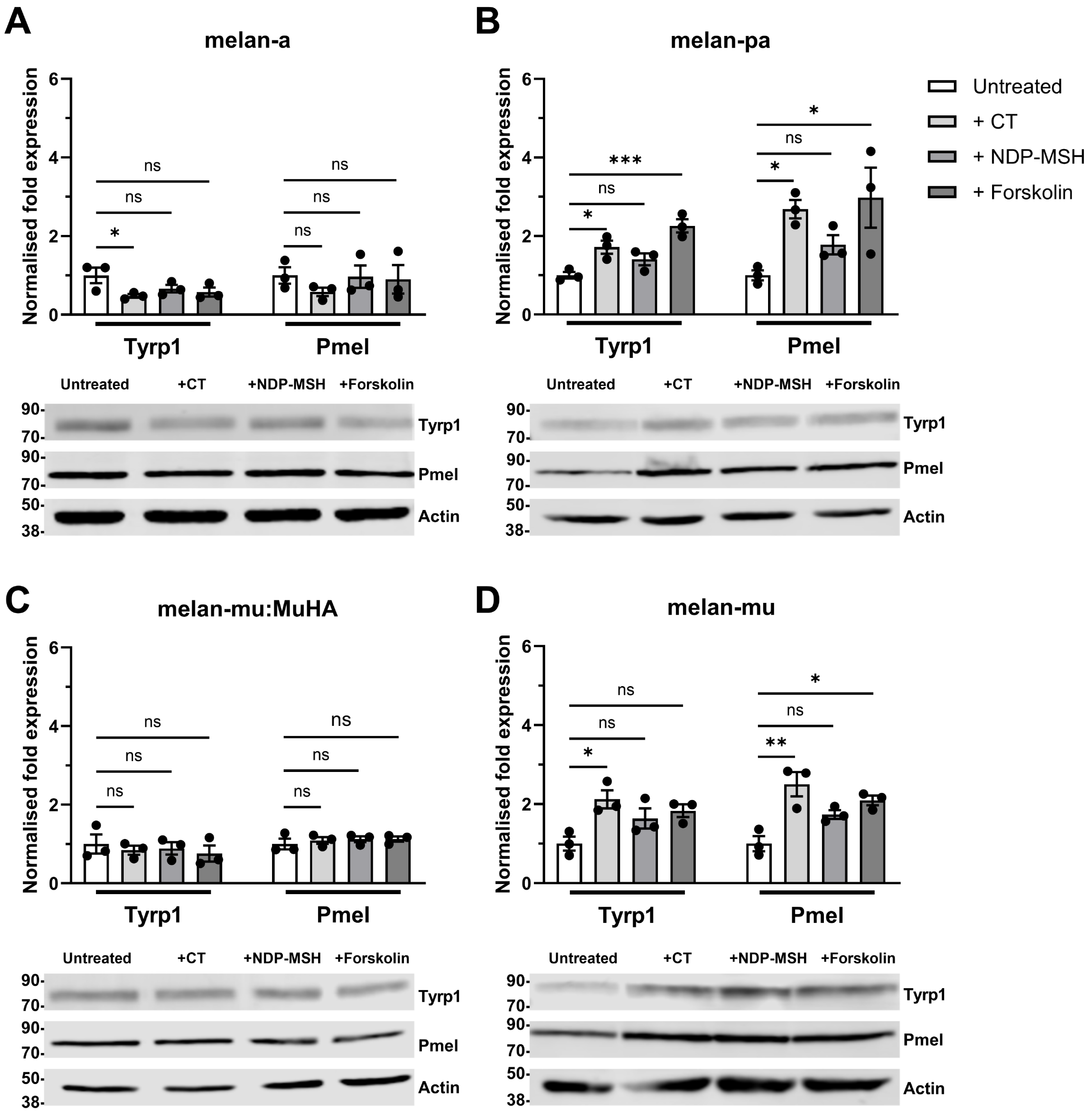


**FIGURE S2. TYRP1 and PMEL protein expression is enhanced upon treatment with cAMP elevating agents in BLOC-1-deficient melanocytes**

BLOC-1 competent (wild-type melan-a, **A**, and BLOC-$1^R$ melan-mu:MuHA, **C**) and BLOC-1-deficient (melan-pa, **B** and melan-mu, **D**) melanocytes were cultured in the absence or presence of the indicated cAMP-elevating agents for 7 days, and then cell lysates were analysed by western blotting for TYRP1, PMEL, and actin as a loading control. Representative blots are shown at bottom, with positions of MW standards (in kDa) shown at the left (for PMEL, the unprocessed P1 form is shown), and quantifications from 3 independent biological replicates are shown at top. Quantitative data are the mean normalised fold expression of TYRP1 or PMEL protein relative to untreated control ± SEM. Statistical significance relative to untreated control was determined by one-way ANOVA with Dunnett's multiple comparison test. Ns, not significant; *, p<0.05; **, p<0.01; ***, p<0.001.

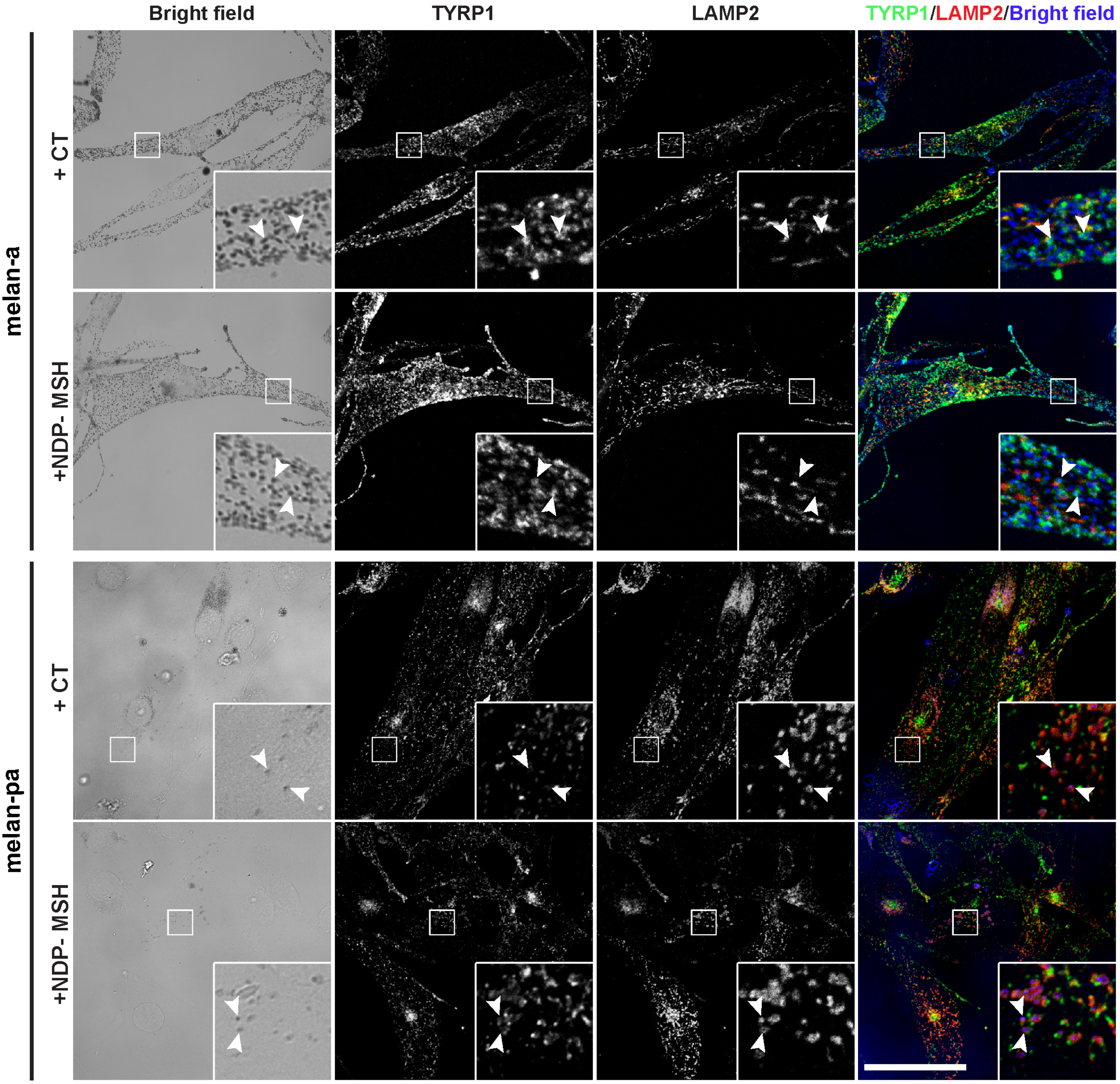


**FIGURE S3. Melanin in BLOC-1-deficient melanocytes resides in LAMP2-positive organelles**

Wild-type melan-a or BLOC-1-deficient melan-pa cells were treated with 200 nM CT or 100 pM NDP-MSH for 7 days, then fixed, immunolabelled, and analysed by confocal IFM and bright field microscopy to determine the localisation of melanin (visible in bright-field images) in relation to melanosomes (TYRP1, green) and late endosomes/lysosomes (LAMP2, red). The bright-field images were inverted and pseudo-coloured blue in the merged image at the right. Insets, boxed regions enlarged 4.5x. Arrowheads show overlap of melanin with TYRP1 but not LAMP2 in melan-a or with LAMP2 but not TYRP1 in melan-pa. Scale bar, 50 µm.

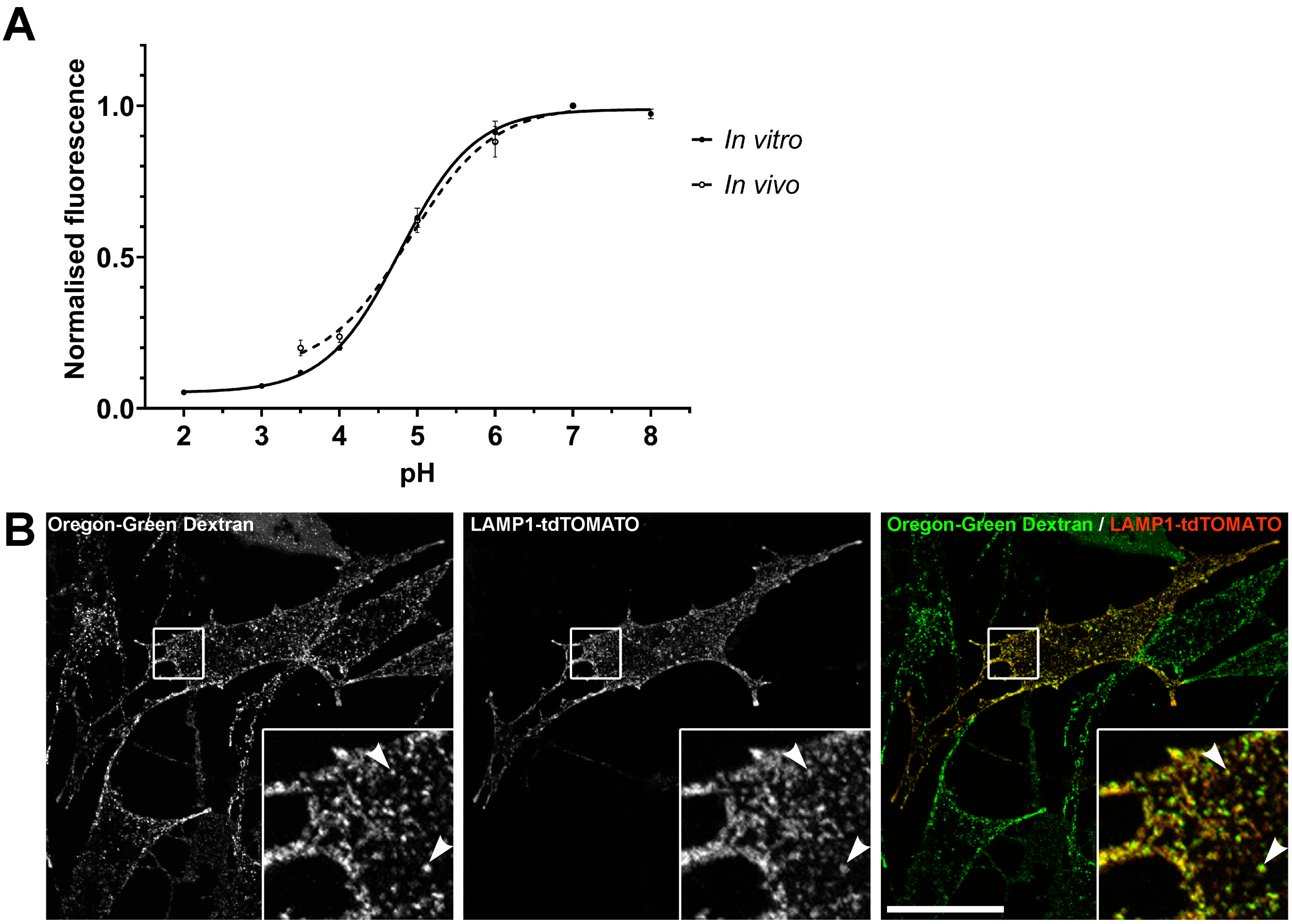


**FIGURE S4. Oregon Green-Dextran is pH sensitive and labels late endosomes and lysosomes.**

**A**. *In vitro* measurements were performed by diluting 50 µg/ml Oregon Green-Dextran in calibration buffers titrated to known pH values (pH 2.0-8.0) and measuring fluorescent intensity using a fluorometer. *In vivo* measurements were obtained using the wild-type melan-a melanocytes that had internalised Oregon Green-Dextran for 16 hours followed by a 2 hour chase period. Cells were then treated with 5 µM monensin and 5 µM nigericin in calibration buffers titrated to known pH values (pH 3.0-8.0) before measuring fluorescent intensity. Both *in vitro* and *in vivo* fluorescence data were normalised to the maximum fluorescence values at pH 7.0 (Oregon Green). A four parameter dose response curve is shown for each condition. Data represent mean ± SEM of 3 independent experiments.

**B.** BLOC-1-deficient melan-pa melanocytes were transiently transfected to express LAMP1-tdTomato and then exposed to Oregon Green-Dextran for 16 hours followed by a 2 hour chase period. Cells were then fixed and imaged by confocal microscopy. Shown is a representative image for each individual label and a merged image at right. Arrowheads point to examples in which labelling for Oregon Green and LAMP1-tdTomato overlap. Scale bar, 50 µm.

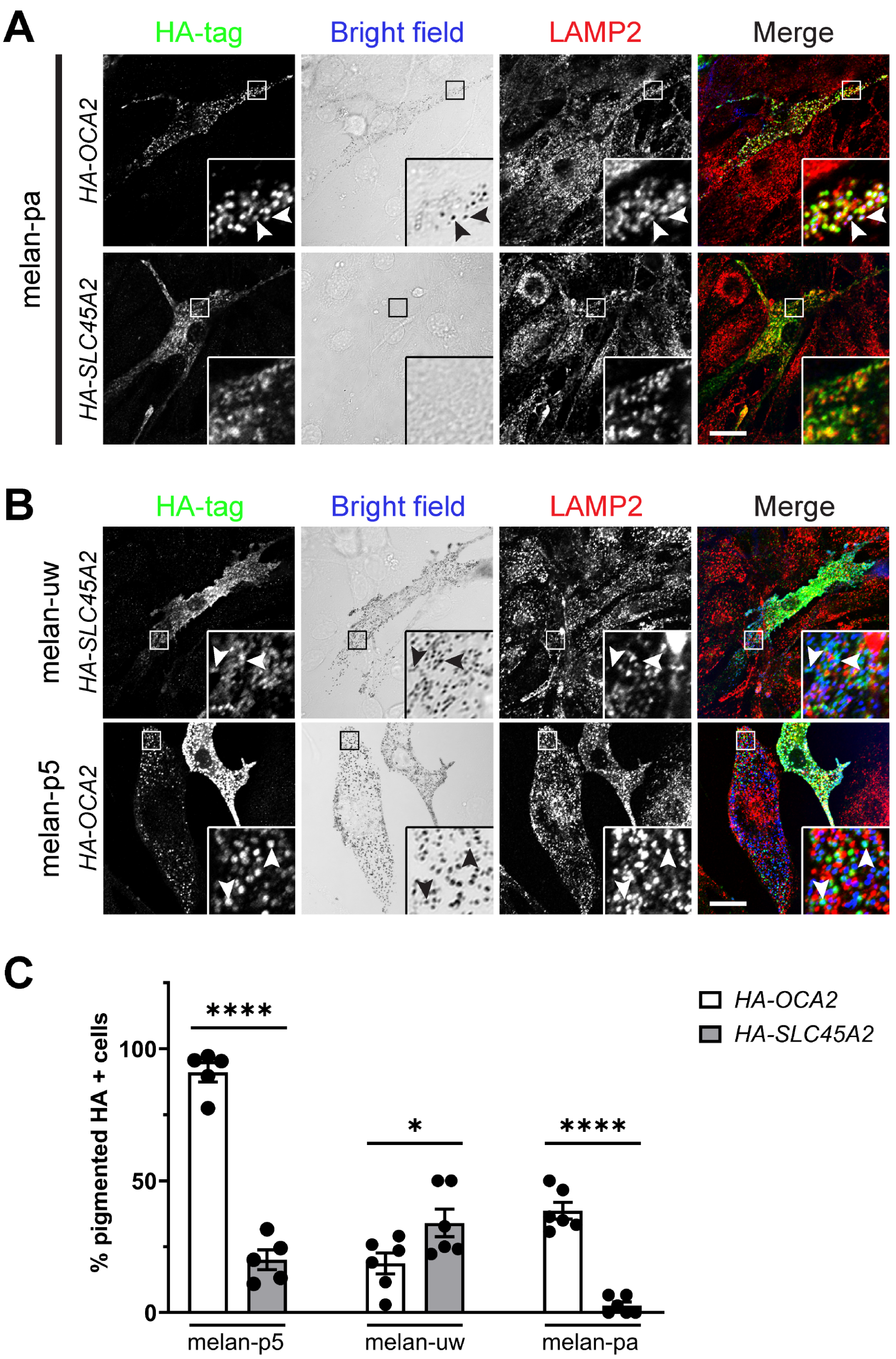


**FIGURE S5. Overexpression of OCA2 in BLOC-1-deficient melanocytes partially restores pigmentation**

BLOC-1-deficient melan-pa melanocytes were transiently transfected with HA-OCA2 or HA-SLC45A2, fixed, immunolabelled for HA and LAMP2 and analysed by confocal IFM and bright-field microscopy to visualise melanin. **A.** Shown are each individual label and a merged image at right which the bright field image was inverted and pseudo-coloured blue. Arrowheads indicate overlap between HA tag, LAMP2 and melanin in cells overexpressing HA-OCA2. Insets of boxed regions are enlarged 4.5x. Scale bar, 20 μm.

**B.** Representative images showing restoration of pigmentation in *Slc45a2*-null melan-uw and *Oca2*-null melan-p5 when transiently transfected with HA-SLC45A2 or HA-OCA2 respectively. Cells were fixed and labelled for HA and LAMP2 and analysed by confocal IFM and bright-field microscopy to visualise melanin. Shown are each individual label and a merged image at right in which the bright field image was inverted and pseudo-coloured blue. Arrowheads indicate overlap between HA tag and melanin in each condition. Insets of boxed regions are enlarged 4.5x. Scale bar, 20 μm.

**C.** Quantification of the number of HA positive melan-uw, melan-p5 and melan-pa cells that were pigmented. Data represent mean ± SEM from 6 independent experiments. Statistical significance relative to the % pigmented SLC45A2 HA positive cells was determined by one-way ANOVA with Dunnett's multiple comparisons test. ns, not-significant; **, $p<0.01$; ****, $p<0.0001$.

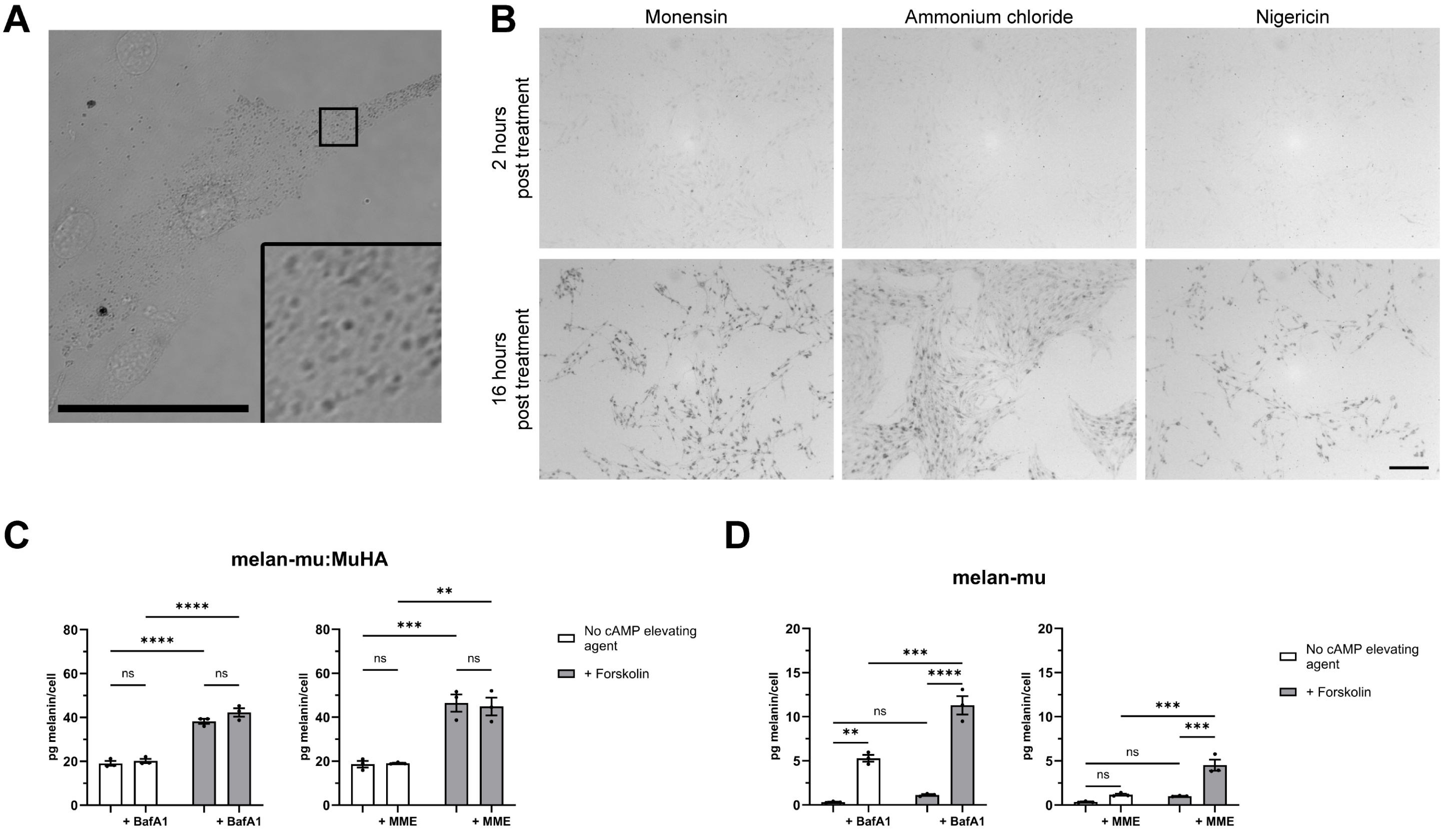


**FIGURE S6. Deacidification agents significantly restore melanin to BLOC-1-deficient melanocytes.**

**A.** Bright field microscopy image of BLOC-1-deficient melan-mu melanocytes treated for 24 hours with 10 mM MME. Inset of boxed region is enlarged 4.5x. Scale bar, 50 μm.

**B.** Bright field microscopy images of BLOC-1-deficient melan-mu melanocytes treated with either 20 mM ammonium chloride, 2 μM monensin, or 2 μM nigericin. Images were taken at 2 and 16 hours post treatment. Scale bar, 200 μm.

**C,D.** Quantification of melanin content of BLOC-$1^R$ (melan-mu:MuHA) **(C)** or BLOC-1-deficient (melan-mu) **(D)** melanocytes cultured for 7 days in the absence or presence of 20 μM forskolin and either 20 nM BafA1 (for final 6 hours) or 10 mM MME (for final 24 hours). Data are from 3 independent experiments and are expressed as mean pg melanin/cell ± SEM. Statistical significance was determined by two-way ANOVA with a post-hoc Tukey test. Ns, not-significant; **, p<0.01; ***, p<0.001; ****, p<0.0001.
